# SPINK1-COL18A1 Crosstalk Shapes Epigenome and Drives Cancer Stemness in Pancreatic Ductal Adenocarcinoma

**DOI:** 10.1101/2025.03.03.641251

**Authors:** Haoyu Tang, Bethsebie Sailo, Xingbo Shang, Paromita Das, Ankit Chhoda, Paulomi Aldo, Marie E. Robert, John Kunstman, Emmanouil Pappou, Laura Wood, Christine A. Iacoubuzio-Donahue, Ralph Hruban, Christopher L. Wolfgang, Linda He, Marie Pfaffl, Olivia Ang Olson, Rolando Garcia-Milian, Mariateresa Mazzetto, Anup Sharma, Andre Levchenko, Nita Ahuja

**Author notes:** These authors contributed equally to this work (Haoyu Tang, Bethsebie Sailo, Xingbo Shang).

## Abstract

Pancreatic ductal adenocarcinoma (PDAC) is a devastating cancer with an increasing incidence and extremely dismal prognosis. Discovery and mechanistic understanding of the genetic and epigenetic drivers of PDAC is therefore of critical importance. Here, we uncover serine protease inhibitor Kazal type 1 (SPINK1) as a putative determinant of clinical progression and demonstrate that it can have a non-canonical function as a regulator of epigenomic states of PDAC cells and associated extensive changes in gene expression. We show that SPINK1 expression, which varies in PDAC, is associated with key aggressive phenotypic cancer states and is correlated with the expression of stemness markers both *in vitro* and *in patient* samples. Mechanistically, our results strongly suggest that SPINK1 acts through a new signaling axis, by interacting with COL18A1 in the Golgi apparatus, promoting endostatin release and eventually inducing extensive histone H3 modifications. These results reveal a new function of SPINK1 in PDAC and suggest a potential therapeutic route to combat PDAC aggressiveness by targeting the SPINK1-COL18A1-endostatin signaling.

## Introduction

Pancreatic ductal adenocarcinoma (PDAC) is one of the most aggressive cancers, characterized by the lowest 5-year survival rate of all major cancer types [1]. At the time of diagnosis, over 75% of patients already present with regionally advanced or metastatic disease, leaving fewer than 20% eligible for curative surgical resection [1–3]. For the majority of patients who cannot undergo surgery, the 5-year survival rate drops to just 2% compared to 11%-34.5% for patients undergoing surgical resection [4, 5]. Due to the limited understanding and therapeutical option, PDAC is projected to become the second leading cause of cancer-related deaths in the United States by 2030 [6]. These realities highlight the critical need for deeper insights into the molecular mechanism of PDAC progression and the development of innovative therapeutic strategies to improve clinical outcomes.

It is increasingly recognized that PDAC progression is driven not only by genetic mutations but also by epigenomic alterations. Our prior research has also demonstrated that PDAC is characterized by extensive epigenomic changes [7]. Distinct PDAC subtypes are revealed with unique epigenomic landscapes, marked by modifications such as H3K4me3 at active promoters, H3K27Ac at active enhancers and promoters, H3K27me3 at Polycomb-repressed regions, and aberrant DNA methylation patterns [8, 9]. Notably, these shifts in epigenomic landscapes could drive transitions between the two primary classifiers–basal and classical subtypes [8]. Some other studies also showed that global epigenetic reprogramming of histone modifications rather than genetic mutations is responsible for the heterogeneity of PDAC tumors [10]. Despite the recognized importance of epigenomic changes contributing to tumor heterogeneity and cellular plasticity that create challenges for effective treatment [11], the mechanisms leading to these changes remain largely unexplored. A few previous studies suggested that mutations or changes in expression of histone modification enzymes are the cause of epigenetic alterations in PDAC [12–18]. However, these results cannot explain all epigenomic changes in PDAC, and further investigation is warranted to discover the underlying mechanisms especially without genetic mutations in epigenetic regulatory genes.

Serine protease inhibitor Kazal type 1 (SPINK1) is initially identified as trypsin inhibitor [19], whose association with pancreatitis is well established [20, 21]. However, its role in PDAC remains controversial [22–27]. While previous studies have shown the role of SPINK1 in several cancers including prostate and hepatocellular cancers [28–30], more research is required to understand the potential contribution of SPINK1 to PDAC progression.

In this study, we examined a cohort of short- and long-term survival PDAC patients (STS and LTS respectively) and discovered a set of genes that potentially contribute to the aggressive clinical outcome. Combined with the analysis of a database with multiple PDAC patient scRNA-seq, this analysis identified SPINK1 as a putative regulator of cancer progression and multiple PDAC associated phenotypes. Further analysis strongly supported the critical role of SPINK1 in modulating cancer cell epigenomics and stemness in PDAC. Subsequent findings led to delineation of a new regulatory pathway, linking SPINK1 to epigenomic alterations through modulation of COL18A1 stability and thus endostatin abundance. Targeting this signaling pathway may offer new therapeutic opportunities for PDAC patients in the future.

## Results

### Integrated analysis of bulk and single-cell transcriptomics from PDAC patients revealed SPINK1 as an important candidate gene for patient survival and tumor development

To identify genes affecting the survival of PDAC patients, we designed an analysis pipeline, combining bulk transcriptomic data from a locally analyzed patient cohort with significant distinction in survival outcome with single-cell data from a published single-cell RNA seq (scRNA-seq) PDAC patient dataset (Fig. 1a).

**Figure 1.**
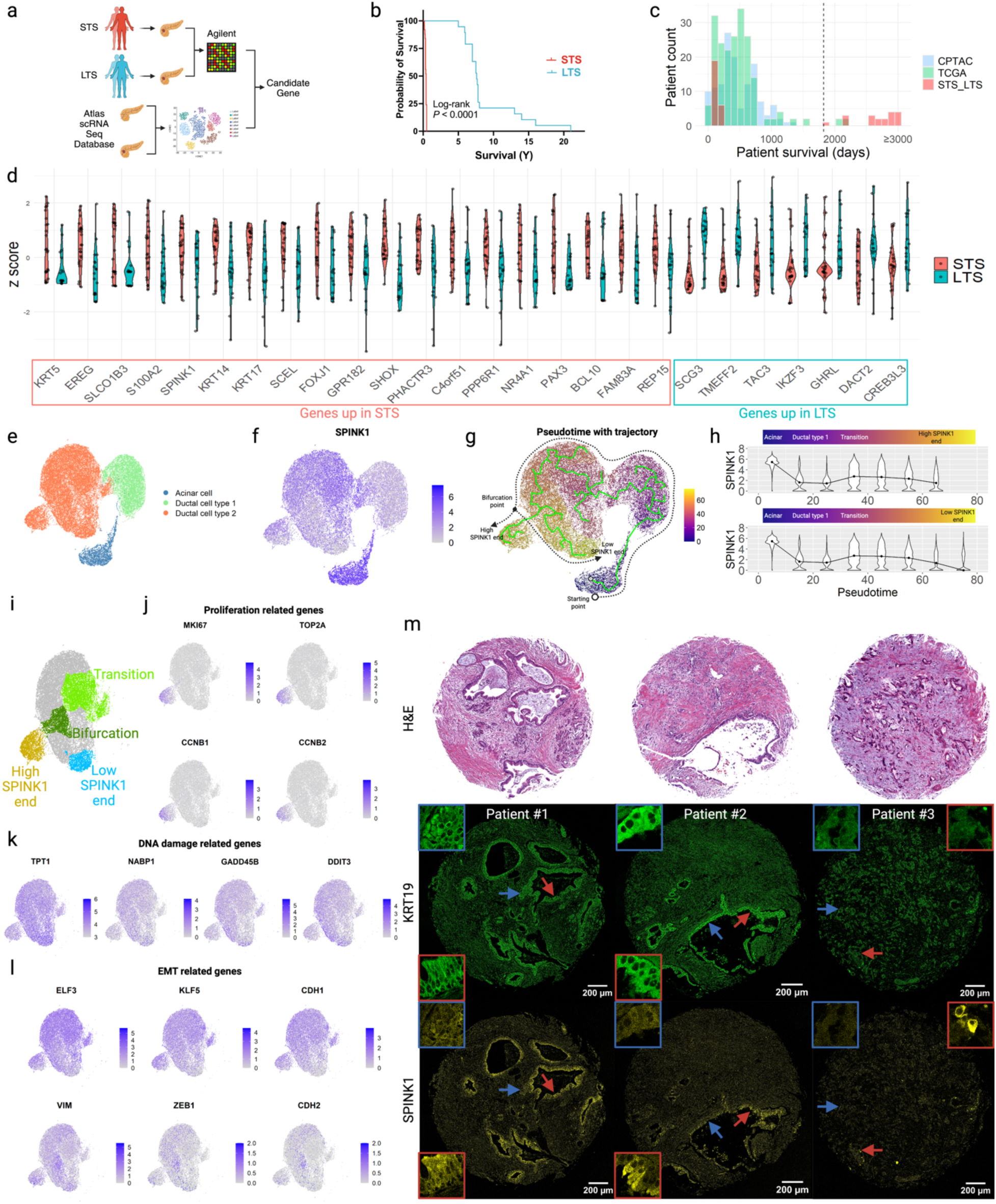
SPINK1 was identified as a gene of interest in PDAC. a) Overall design of the integrated analysis of bulk and single-cell transcriptome in PDAC patients. b) Survival curves of STS and LTS in the patient cohort used for bulk transcriptome analysis. c) Histogram comparison of patient survival from TCGA, CPTAC and the STS-LTS cohort used in this study. The vertical black dashed line marked survival of 5 years (1825 days). d) Violin plots of z scores of the 26 short list genes identified from the bulk transcriptome analysis of the STS-LTS cohort (|log2FC| ≥ 1.5 and FDR ≤ 0.25). e) UMAP visualization of acinar and ductal cells from the reference dataset integrated by Harmony across different patients. f) Visualization of SPINK1 level in e). g) Visualization of trajectory and pseudotime inferred by Monocle3 in e). (The trajectory drawn in the solid green line is the direct output from Monocle3 whereas the black dashed line is drawn manually based on the solid green line for illustration purposes.) h) Violin and line plots of SPINK1 level along the trajectories towards high SPINK1 end (upper panel) and low SPINK1 end (lower panel). Cells were binned in pseudotime with a bin-size of 10. The medians of the violin plots were marked and joined by lines. i) UMAP visualization of subclusters defined in ductal cell type 2. j) UMAP visualization of proliferation and k) DNA damage related genes in ductal cell type 2. l) UMAP visualization of EMT related genes in ductal cell type 2. m) Exemplary PDAC FFPE sections with different overall SPINK1 protein levels. Zoomed views of ROIs pointed by the red and blue arrows are shown in the inset with the corresponding color. ROIs pointed by the red (blue) arrows have relatively high (low) SPINK1 levels within each patient. Patient #1, #2, and #3 corresponds to PA-5, PA-2, PB-7 in Fig. S3.

More specifically, we first curated a unique patient cohort consisting of 25 STS (survival ≤1 year) and 19 LTS (survival ≥5 years) (Fig.1b), which were age, gender, and tumor location matched (Table S1). Due to the high lethality of PDAC, currently available patient cohorts such as those from TCGA [31] and CPTAC [32] have none or very few LTS representatives. By contrast, our unique cohort contains a significantly larger number of LTS with stringent selection criteria (Fig. 1c), which renders it possible to uncover previously unidentified gene expression patterns associated with patient survival. In total, a shortlist of 26 differentially expressed genes (DEGs) with |log2FC| ≥ 1.5 and false discovery rate (FDR) ≤ 0.25 were identified by comparing the bulk transcriptomic data of STS with LTS (Fig. 1d). Notably, three known predictors of poor PDAC prognosis, KRT5, KRT17 and S100A2 [33, 34], were present in our shortlist as genes upregulated in STS, which added to the validity of our approach.

To further dissect the functional role of these short-listed genes in the tumor microenvironment, we examined a reference PDAC scRNA-seq dataset combining a large number of patient and normal donor samples to study the transcription level of each gene in our shortlist at single-cell resolution [35]. After correcting several key errors in the original reference dataset (see Methods and Fig.S1a-1b for details), the distribution of each gene from the short list across different cell types was analyzed (Fig.S1c-1d). Similar to previous observations [36], pancreatic ductal cells grouped into two different clusters (Fig. 1e), one of which (ductal type 1 cells) was normal-like, as evidenced by the presence in normal control samples, whereas the other (ductal type 2 cells) was likely the cluster of cancer cells, as evidenced by the dominant presence in tumor samples and low trypsin but high KRT19 expression (Fig. S2a-2b). This distribution revealed a striking elevation of SPINK1 expression in both acinar (as expected based on the anti-trypsin function in the normal pancreas) and, importantly, in ductal type 2 but not type 1 cells (Fig. 1f). We also observed highly heterogeneous SPINK1 level in only ductal type 2 cells. Furthermore, the expression of SPINK1 in the putative PDAC cells (ductal type 2) was markedly higher than the other shortlisted gene, suggesting its potential significance in cancer cells rather than in the tumor microenvironment.

To understand how this variability of SPINK1 aligns with tumor progression, a pseudotime trajectory was inferred by Monocle3 [37–39] with acinar cells as the starting point based on a common theory of acinar-to-ductal metaplasia (ADM) in PDAC carcinogenesis [40]. The trajectory traversed acinar, ductal type 1 and ductal type 2 cell clusters, and bifurcated, leading to two terminal points (‘ends’) in ductal type 2 cluster (Fig.1g). Based on the finding from Fig.1f, one trajectory end was found with significantly higher SPINK1 level compared to the other (coverage of two trajectories with different end was shown in Fig. S2c). We next plotted the SPINK1 level along the trajectories of two different ends (Fig. 1h). We observed that SPINK1 expression dropped during the transition from acinar cell cluster to normal-like ductal cell type 1 cluster and subsequently strongly increased when transiting into tumor-like ductal cell type 2 cluster. After the increase in this ‘transition zone’, the SPINK1 level partially decreased along the trajectory which then underwent a major bifurcation with the trajectory ends displaying high and low SPINK1 expression as indicated above.

To dissect the changes in the transcriptomic and associated phenotypic states along the inferred trajectory, based on the above observations, we isolated within the ductal cell type 2 cluster four different subclusters (*i*. transition zone with elevated SPINK1 expression, *ii.* bifurcation zone, *iii.* high SPINK1 end and *iv*. low SPINK1 end) based on their position in the trajectory (Fig. 1i). We first focused on the difference between the high SPINK1 end and low SPINK1 end subclusters, and found relatively high expression of proliferation related genes (MKI67, TOP2A, CCNB1, CCNB2 [36]) in cells of high SPINK1 end subcluster, indicating a proliferative state, while high expression of DNA damage related genes (TPT1 [41], NABP1 [42], GADD45B [43], DDIT3 [44]), indicating a state with the stress of DNA damage, was found in the low SPINK1 end subcluster (Fig.1j-k). In addition, the transition subcluster compared to especially the bifurcation subcluster largely overlapped with the group of cells displaying high epithelial related markers (ELF3, KLF5 and CDH1 [45, 46]) and/or lower epithelial-mesenchymal transition (EMT) markers (VIM [47], ZEB1 [48] and CDH2 [46]) (Fig. 1l). These findings indicated that SPINK1 expression elevates in cells that, after transitioning to the cancerous state, display active epithelial, EMT and proliferation phenotypes. Conversely, its expression decreases in cells that adopts a DNA damage stress response phenotype.

To verify whether SPINK1 heterogeneity observed in the scRNA-seq reference dataset can also be observed at protein level in patients, we co-stained SPINK1 and epithelial marker KRT19 in a tissue microarray (TMA) (Fig. S3). Tumor regions were determined by both KRT19 immuno-staining and corresponding H&E from adjacent slices. In agreement with the findings from the scRNA-seq dataset, substantial heterogeneity in SPINK1 protein levels was observed not only across different patients but also in different tumor regions within individual patient or tumor ductal cells within individual tumor regions (representative examples shown in Fig. 1m).

Overall, our analysis implicated SPINK1 as one of the genes associated with STS in PDAC patients, whose expression is particularly elevated in ductal type 2 cells, corresponding to PDAC cells. Furthermore, the trajectory analysis showed that the expression of SPINK1 is associated with diverse cancer phenotypes and is particularly upregulated during the transition to carcinogenesis. These findings suggested that SPINK1 might modulate multiple transcriptional programs in highly plastic pancreatic cancer cells.

### SPINK1 knockout induces substantial and consensual changes in the transcriptome and chromatin accessibility in PDAC

Following the finding that SPINK1 is a gene potentially related to patient outcomes and associated with diverse phenotypic outcomes, we probed SPINK1 function in a battery of PDAC cell lines. Considering the high heterogeneity of SPINK1 observed in scRNA-seq, we postulated that such heterogeneity might also exist across different PDAC cell lines and that *in vitro* study should be carried in multiple cell lines spanning different levels of SPINK1 for the validity of the results. Therefore, the cell models were selected based on SPINK1 mRNA levels as reported in the Cancer Cell Line Encyclopedia (CCLE) database [49] (Fig. S4a). In five cell lines, we confirmed that gene expression levels indeed correlated with SPINK1 protein levels (Fig. S4b-4c). Based on the results, MiaPaCa2 and PANC-1 cell lines were chosen as the representative PDAC cell lines with undetectable and low SPINK1 expression respectively, while BxPC-3 and HPAF-II cell lines were chosen as the representative cell lines with high SPINK1 expression for the subsequent *in vitro* experiments.

We first knocked out SPINK1 in PANC-1 and BxPC-3 cell lines using CRISPR-Cas9 gene targeting (Fig. S4d-S4e) and found that the loss of SPINK1 led to a significant decrease in cell proliferation, migration, and invasion ability (Fig. S4h-4l), which supported a role of SPINK1 in regulation of diverse PDAC-associated phenotypes. To gain further information on gene regulation downstream of SPINK1, we next applied bulk RNA-seq to SPINK1 knockout (sgSPINK1) and scramble control (sgCtrl) PANC-1 cells. We indeed observed wide-ranging changes in the transcriptome (2163 DEGs) after the knockout of SPINK1 (Fig. 2a), suggesting involvement of mechanisms that may not depend on a single transcription factor or program. Changes in chromatin accessibility are an important upstream regulator of transcription, frequently occurring in cancers including PDAC and affecting the expression of large numbers of genes. To examine if the significant shift in the transcriptome in sgSPINK1 cells was due to changes in the chromatin accessibility, bulk ATAC-seq was also performed and revealed a substantial change in chromatin accessibility after SPINK1 knockout (Fig. 2b-2c). By intersecting the results from RNA-seq and ATAC-seq, 1862 genes with significant changes in both transcription level (adjusted p-value ≤0.05) and chromatin accessibility (FDR ≤0.05) were identified. In alignment with a strong coupling between chromatin accessibility and transcription, 873 and 655 genes were consensually upregulated and downregulated respectively out of these 1862 genes (representing 82% of all genes involved; Fig. 2d), which strongly indicated that SPINK1 associated gene expressions are indeed associated with epigenetic alterations.

**Figure 2.**
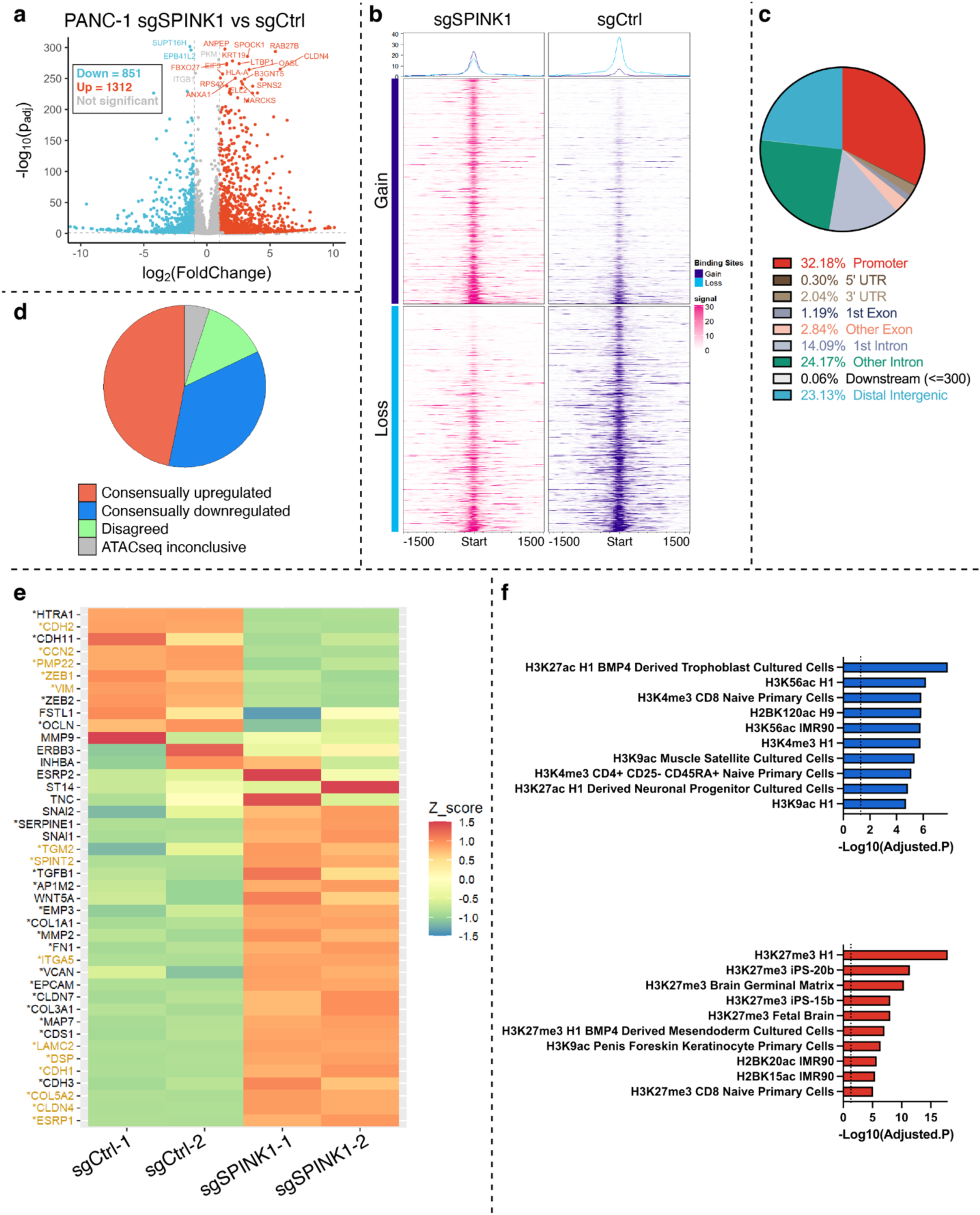
SPINK1 knockout induced significant and consensual changes in the transcriptome and chromatin accessibility in PANC-1 cells. a) Volcano plot from the bulk RNA-seq (n = 2 sgSCR, n =2 sgSPINK1). Differentially expressed genes (DEGs) with adjusted p value ≤ 0.05 and log2 fold change ≤ -1 or ≥ 1 were highlighted in blue and red respectively. b) Overview of sites with gain and loss in chromatin accessibility identified from ATAC-seq (n = 2 sgSCR, n =2 sgSPINK1). c) Pie chart of different genomic locations annotated to differentially accessible ATAC-seq peaks. d) Pie chart of genes identified as differentially regulated in both RNA-seq and ATAC-seq. If one gene simultaneously contains sites with upregulated and downregulated accessibility, it is labeled as ATAC-seq inconclusive. If one gene has accessibility and transcription level changing in the opposite direction, it is labeled as disagreed. e) Heatmap of Z score of EMT related genes calculated from normalized counts from RNA-seq. Genes with differential transcription level (adjusted p value ≤ 0.05) were labelled with asterisk sign. Genes with differential accessibility (FDR ≤ 0.05) were highlighted in yellow. f) Enrichment result from Enrichr using the Epigenomics Roadmap HM ChIP-seq database for consensually downregulated (upper panel) and upregulated (lower panel) genes. The lengths of the bars corresond to the values of -log10(adjusted p value), and the dash lines indicate the cut-off (adjusted p = 0.05).

We then focused on these consensually up/downregulated genes because they were more likely to be the direct targets of SPINK1-induced chromatin alterations. Particularly, we found that multiple mesenchymal markers (CDH2, VIM, ZEB1) and epithelial markers (CDH1) were consensually downregulated and upregulated after SPINK1 knockout respectively (Fig. 2e), suggesting a potential role of SPINK1 in epigenetic control of EMT. Enrichment analysis with Enrichr [50] showed that two activating histone modifications [51–53] H3K4Me3 and H3K27Ac were enriched for consensually downregulated genes (Fig. 2f upper panel), whereas H3K27Me3, a repressing modification [51], was enriched for consensually upregulated genes (Fig. 2f lower panel). This implied that SPINK1 might exert an epigenetic control on the transcriptome of PANC-1 cells by regulating histone modifications.

Overall, these data suggested that SPINK1 might indeed control diverse, PDAC-associated transcriptional programs through regulation of chromatin accessibility and epigenomic alterations.

### SPINK1 modulates the epigenome of PDAC

To verify the integrated results from ATAC-seq and RNA-seq analysis, western blot was utilized to investigate the effect of SPINK1 on EMT in PDAC cell lines. We chose four EMT related genes: CDH1 (E-Cadherin), CDH2 (N-Cadherin), ZEB1 and VIM (Vimentin) because of their well-known role in EMT and significant alterations in both transcript levels and chromatin accessibility in PANC-1 cell line after SPINK1 knockout (Fig. 2e). In agreement with the sequencing results, SPINK1 knockout PANC-1 cells indeed showed increasing protein level of epithelial marker (E-Cadherin) and decreasing protein levels of mesenchymal markers (N-Cadherin, ZEB1, and Vimentin), indicating a decrease in EMT (Fig. 3a-3b). However, in another PDAC cell line, BxPC-3, immunoblot analysis showed no significant changes in these four EMT related genes after SPINK1 knockout. Additionally, after lentiviral mediated overexpression of SPINK1 in all four selected cell lines (Fig. S4f-S4g), only PANC-1 was found to have corresponding changes in these four EMT related genes (Fig. 3c and 3d). These findings suggested that SPINK1 alone may not be sufficient for controlling EMT and that some factors present in PANC-1, but absent in the other three cell lines, are required to mediate SPINK1 regulation of EMT in PDAC.

**Figure 3.**
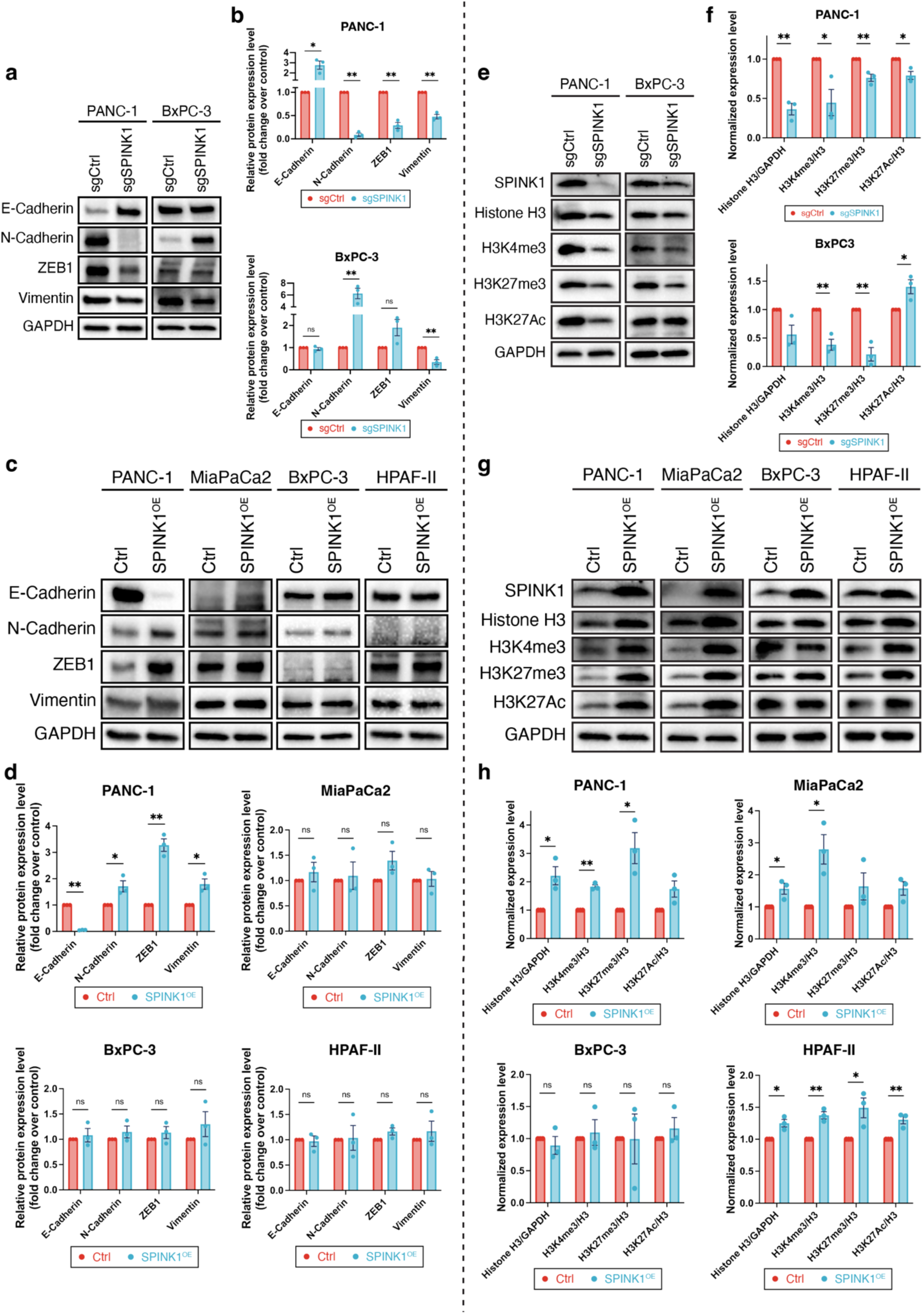
SPINK1 was not the direct regulator of epithelial-to-mesenchymal transition (EMT) but an essential modulator of the epigenome in PDAC. (a) Western blot showed the effect of knock-out of SPINK1 on EMT in PANC-1 and BxPC-3 cells (quantification in 3b). (c) Western blot showed the effect of over-expression of SPINK1 on EMT in four PDAC cell lines (quantification shown in 3d). (e) Western blot showed the effect of SPINK1 knock-out on histone H3 and its modifications in PANC-1 and BxPC-3 cells (quantification in 3f). (g) Western blot showed the effect of SPINK1 over-expression on histone H3 and its modifications in PDAC cell lines (quantification in 3h).

We then assessed the effect of SPINK1 perturbation on the three histone modifications (H3K4me3, H3K27me3, and H3K27Ac) suggested by the RNA-seq and ATAC-seq data (Fig. 2f). We observed that the knockout of SPINK1 led to a significant decrease in histone H3 protein as well as its modifications (H3K4me3 and H3K27me3) in both PANC-1 and BxPC-3 cell lines (Fig. 3e-3f). Correspondingly, overexpression of SPINK1 resulted in a significant increase in histone H3 protein and two histone modifications (H3K4me3 and H3K27me3) in all selected PDAC cell lines except for BxPC-3 (Fig. 3g-3h). Since BxPC-3 cells showed high baseline expression of SPINK1 (Fig.S4b-4c), we reasoned that the absence of SPINK1 overexpression to modulate epigenomics was due to the saturation of SPINK1 signaling in wild-type BxPC-3. These observations, which shows the regulatory effect of SPINK1 on histone modifications remained consistent across different PDAC cell lines with differing SPINK1 baseline levels, indicates that SPINK1 is indeed an important modulator of the epigenome in PDAC.

### SPINK1 expression is correlated with cancer stemness and is negatively regulated by EMT in PDAC

After establishing the link between SPINK1 and histone modifications, we proceed to explore the genes and functions regulated by this connection. Cancer stem cells have been widely implicated in controlling tumor heterogeneity and contributing to tumor recurrence and eventually poor clinical outcomes [54]. Since SPINK1 was found to be associated with shorter survival, epigenomic alteration, diverse transcriptomic programs and associated phenotypes i.e., plasticity of cancer cells, we investigated if there is a correlation between SPINK1 expression and cancer stemness in PDAC. When examining the enrichment results from Enrichr for consensually downregulated genes in PANC-1 SPINK1 KO cells, we noticed stemness related terms were top-rated in results based on three different databases (Fig. 4a). Moreover, the leading-edge genes for stem cell related terms largely overlapped with the leading-edge genes for histone modifications (H3K4Me3 and H3K27Ac) from the previous findings (Fig. 4b), indicating SPINK1 might modulate stemness through regulating epigenomics. Given that cancer stem cells (CSC) constitute an important source of tumor heterogeneity and contribute to tumor recurrence and eventually poor clinical outcome [54], we hypothesized that SPINK1 negatively impacted patient survival by increasing the stemness of PDAC tumor cells.

**Figure 4.**
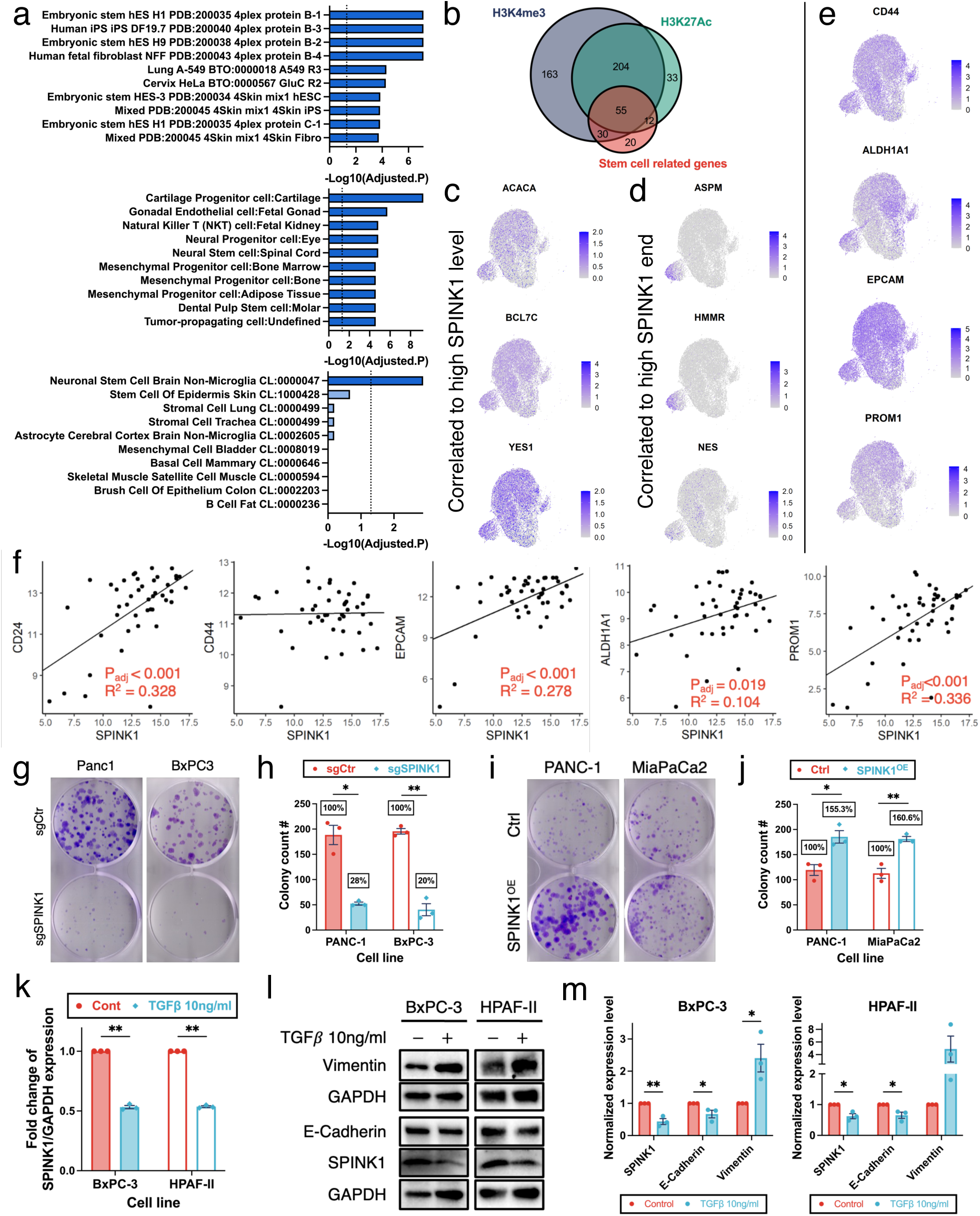
SPINK1 regulates PDAC stemness and plasticity. (a) Enrichment of consensually down-regulated genes based on ProteomicsDB 2020 (upper), CellMarker Augmented 2021 (middle), Tabula Muris (lower) databases. The dash lines indicate the cut-off (adjusted p = 0.05). (b) Venn diagram showed the overlapping of stem cell related genes from Fig. 4a with H3K4me3 and H3K27Ac regulated genes from Fig. 2f. (c) UMAP projections of representative stem cell related genes correlated to high SPINK1 level. (d) UMAP projections of representative stem cell related genes exclusively enriched in the high SPINK1 end subcluster (subcluster iii). (e) UMAP plots of representative cancer stemness markers in Ductal cell type 2 cluster. (f) The correlation plots of representative cancer stemness marker genes in our initial STS_LTS patient cohort. (g) Colony formation assays of SPINK1 knock-out in PANC-1 and BxPC3 cells (quantification in 4h). (i) Colony formation assay of SPINK1 over-expression in PANC-1 and MiaPaCa2 cells (quantification in 4j) (k) qPCR showed the transcriptional level of SPINK1 in BxPC-3 and HPAF-II cells with or without TGF*β* treatment. Data are presented as the mean ± s.e.m., analyzed using a two-tailed unpaired Student’s *t*-test, with *n* = 3 independent experiments per cell type. (l) Western blot revealed the effect of TGF*β*-induced EMT on SPINK1 protein levels in BxPC-3 and HPAF-II cells (quantification shown in 4m).

In search of support for this hypothesis, we next explored the relationship between SPINK1 expression and cancer stemness in PDAC patient data. The scRNA-seq dataset showed stemness related genes with a type of expression pattern undergoing a similar bifurcation as SPINK1 with low (high) expression in low (high) SPINK1 end (Fig. 4c). However, another distinctive group of stemness related genes was almost exclusively enriched in the high SPINK1 end subcluster (i.e., in cells displaying high proliferation) (Fig. 4d). Additionally, the expression of four commonly used CSC markers, EPCAM, CD44, ALDH1A1, and PROM1 (CD133), was enriched in cells displaying high expression of SPINK1, with the latter three showing particularly high expression in the transition and high SPINK1 end subclusters (Fig. 4e). Lastly, reexamination of comparison between STS versus LTS patient cohorts revealed that the expression of ALDH1A1, EPCAM, PROM1 together with another widely accepted PDAC CSC marker CD24 were all significantly correlated to the SPINK1 expression in PDAC patients (Fig. 4f).

Since tumor initiation is a critical characteristic of CSCs [55], we performed colony formation assays confirming that SPINK1 KO significantly reduced the number of colonies formed by PANC-1 and Bxpc3 cells in agreement with our hypothesis (Fig. 4g-4h). In contrast, SPINK1 overexpression significantly increased the number of colonies formed by PANC-1 and MiaPaCa2 cell lines (Fig. 4i-4j). Altogether, these results from both *in vitro* and *in patient* analysis suggest that there is a positive correlation between SPINK1 and PDAC stemness likely mediated by SPINK1 regulated histone modifications.

Stemness has previously been associated with the ability to adopt diverse phenotypic states, including EMT, provided that corresponding cues exist in the cellular micro-environment. This transition may correspond to the EMT sub-cluster observed in the patient cells and SPINK1-dependent phenotype in PANC1 cells described above (Fig. 1l). To further investigate the interplay between stemness and EMT in PDAC cell lines, we treated BxPC-3 and HPAF-II wildtype cells with transforming growth factor-β (TGFβ), a common EMT inducer, and found significant decrease in the transcription of SPINK1 by qPCR (Fig.4k). Further, the decrease of SPINK1 protein level in both cell lines after TGFβ-induced EMT was subsequently confirmed by Western blot (Fig. 4l-4m). This indicates that EMT process may be partially enabled by SPINK1, but it also causes a decrease in SPINK1 level through regulating its transcription. This mechanism may explain the prior observation of SPINK1 decreasing in the cells undergoing EMT along the trajectory bifurcating to distinct SPINK1 ends (Fig. 1h).

Overall, our results from both *in vitro* and tissue samples suggest that there is a positive correlation between SPINK1 and PDAC stemness, possibly through SPINK1’s modulation of the epigenome. Further, the EMT process during cancer progression may in turn decrease SPINK1 expression in tumor cells.

### COL18A1 is an essential interacting partner of SPINK1 responsible for epigenomic changes

After exploring the association between SPINK1 and cancer stemness through histone modifications in PDAC, we set out to investigate the mechanism by which SPINK1 can control these epigenomic alterations. A prior study by Ozaki et al. reported that recombinant SPINK1 (rSPINK1) bound to and activated EGFR in PDAC cells under serum free condition [56]. Since all preceding observations of SPINK1’s effects on histone modifications in our study were performed in serum-complemented medium, we next investigated whether SPINK1 worked as an EGFR agonist also in the medium supplemented with 10% FBS. Although the positive control EGF could clearly increase the phosphorylation of EGFR, indicating EGFRs were not saturated by complete medium, exposure to high dose of rSPINK1 led to no detectable activation on EGFR in different PDAC cell lines (Fig. S5a & S5b). We therefore hypothesized that there is an alternative signaling pathway of SPINK1 without the involvement of EGFR in PDAC.

To research alternative mechanisms of SPINK1 mediated epigenetic control, we first relied on the conventional immunoprecipitation-mass spectrometry (IP-MS) approach but were unable to reproducibly identify any interaction partner. We reasoned that the interaction between SPINK1 and the unknown partner(s) could be too weak to withstand the harsh washing condition during IP and then switched to the proximity dependent biotinylation method. To minimize the impact on SPINK1 from the connected biotinylation enzyme, we fused UltraID, a small and highly reactive biotinylation enzyme, recently developed by Kubitz et al. [57], to SPINK1 (SPINK1-UltraID) and used UltraID fused to the signaling peptide of SPINK1 (Ultra Ctrl) as the negative control in PANC-1 cells. During preliminary small-scale testing using HRP-conjugated streptavidin, we observed a biotinylated band slightly above 150kDa showing a distinctly stronger signal in lysate from SPINK1-UltraID (Fig. S5c and S5d). We then performed pull-down using streptavidin beads at larger scale and analyzed the short peptides mixture obtained from on-bead digestion using MS (Fig. 5a). We determined a shortlist of possible interaction partners according to 5 criteria—molecular weight between 150 and 200 kDa, ratio of iQAB value of SPINK1-UltraID to that of Ultra Ctrl larger than 2, no protein grouping ambiguity, total spectrum count more than 1, and absolute log2FC value from prior RNA-seq of SPINK1 knockout less than 2 (Fig. 5b). Within the shortlist, COL18A1 was shown as the top candidate with a hundred times higher iQAB signal compared to the other hits. The western blot analysis also supported that more COL18A1 was indeed specifically pulled down from SPINK1-UltraID cells using streptavidin beads in comparison to the negative control (Fig. 5c). Furthermore, when DSP crosslinker was added to stabilize the complex, COL18A1 could be co-immunoprecipitated (co-IP) with SPINK1 in PANC-1 cells overexpressing SPINK1-ALFA using nanobodies highly specific to the ALFA tag (Fig. 5d). We confirmed that the co-IP of COL18A1 was specific to SPINK1 and could not be found in the control cells overexpressing only ALFA tag (Fig. 5e). We further showed that SPINK1 colocalized with COL18A1 in the Golgi apparatus in both PANC-1 and MiaPaCa2 cells based on immunostaining (Fig. 5f). This result matched with the subcellular protein fractionation findings suggesting that both SPINK1 and COL18A1 were most abundant in membrane-bound organelles (Fig. S5e). We therefore proposed that COL18A1 was an interacting partner of SPINK1 inside the Golgi apparatus in PDAC cells.

**Figure 5.**
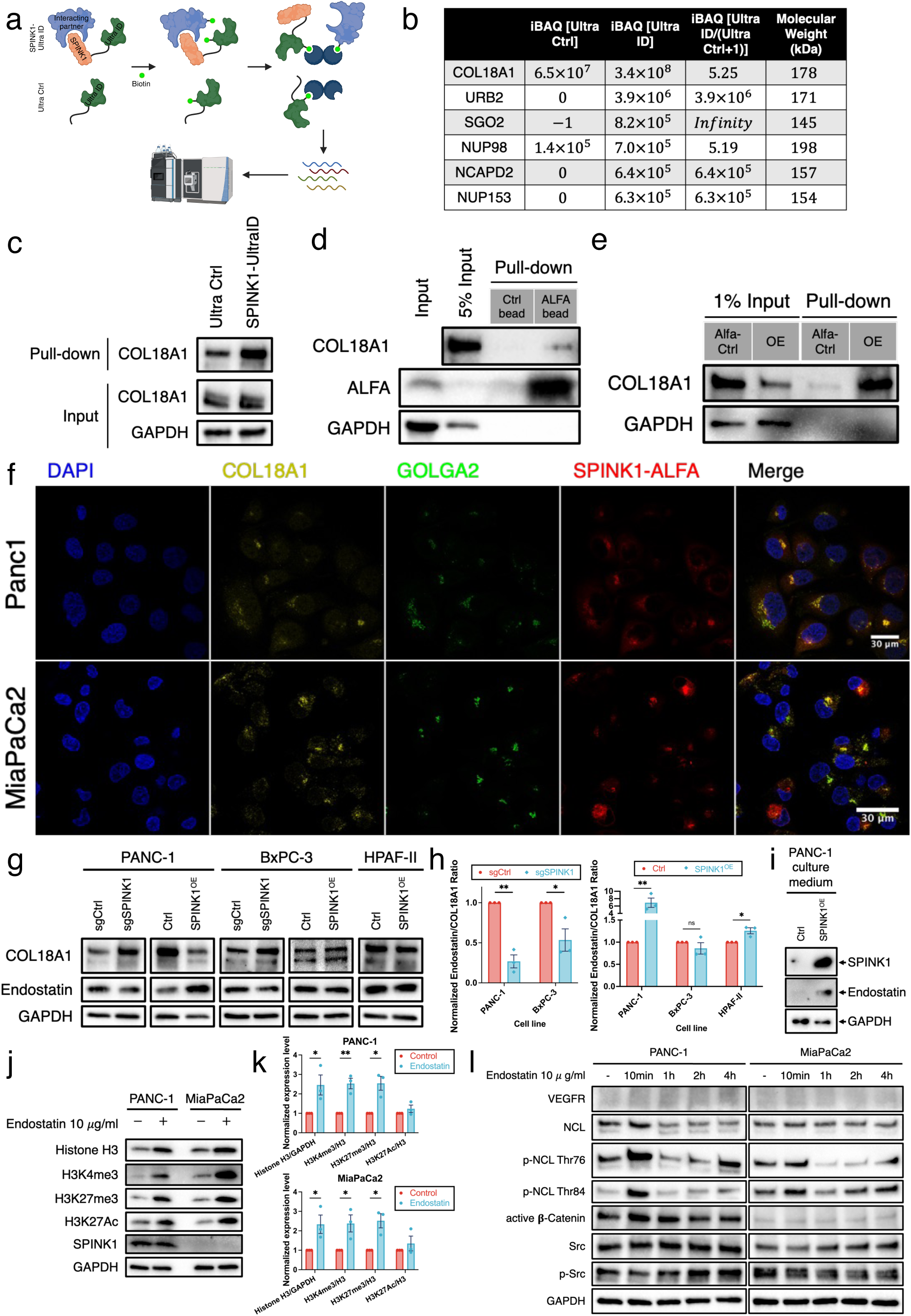
SPINK1 regulates epigenomics through interacting with COL18A1 in PDAC. (a) A schematic shows the experiment design of immunoprecipitation with proximity labeling by Ultra ID enzyme to identify interacting partner(s) of SPINK1. (b) The table of candidate partners of SPINK1 based on the mass spectrometry results after on-bead digestion. (c) Western blot showed the IP of biotinylated COL18A1 in the PANC-1 Ultra ID cell system using streptavidin beads. (d) Western blot showed the co-IP of COL18A1 using SPINK1^OE^ PANC-1 cells with crosslinker DSP and Ctrl beads (bead surface with IgG) or ALFA beads (bead surface with nanobodies against ALFA tag). (e) Western blot showed the co-IP of COL18A1 in Ctrl versus SPINK1^OE^ PANC-1 cells with the crosslinker and ALFA beads. (f) Confocal microscopy showed the immunostaining of SPINK1, COL18A1 and Golgi apparatus marker GOLGA2 in PANC-1 and MiaPaCa2 cells. (g) Western blot revealed the effect of SPINK1 knock-out or over-expression on endostatin/COL18A1 ratios in PDAC cell lines (quantification in 5h). (i) Western blot revealed the released SPINK1 and endostatin from SPINK1^OE^ PANC-1 cells to the culture medium. (j) Western blot revealed the effect of recombinant endostatin on the histone H3 changes in PANC-1 and MiaPaCa2 WT cells after 1-day treatment (quantification in 5k). (l) Western blot revealed the effect of endostatin on potential downstream targets in PANC-1 and MiaPaCa2 cells (quantification in S5i).

It is widely known that the C-terminal domain of COL18A1 can be cleaved to produce a small protein known as endostatin [58]. We thus next investigated whether SPINK1’s interaction with COL18A1 would affect the production of this bioactive peptide. We found that the knockout of SPINK1 decreased the ratio of endostatin versus COL18A1 in both PANC-1 and BxPC-3 cells, whereas overexpression of SPINK1 led to a significant increase in the ratio of endostatin versus COL18A1 in all PDAC cell lines except for BxPC-3 (Fig. 5g-5h). Notably, the baseline ratios of endostatin versus COL18A1 were also significantly higher in PDAC cell lines with high SPINK1 expression than those with low or undetectable SPINK1 expression (Fig. S5f & S5g). These results suggested that the interaction between SPINK1 and COL18A1 promoted the cleavage of COL18A1 to generate more endostatin. To be noticed, the partial deletion experiment demonstrated that the deletion of residues 25-34 showed the strongest diminishing effect on SPINK1-induced histone H3 alterations (Fig. S5h), which indicated the key amino acid in this function might be different from the lysine residue at position 41 responsible for trypsin inhibition. By measuring the concentrated conditional medium of PANC-1 cells, we confirmed the extracellular release of endostatin, which was enhanced by elevated SPINK1 expression (Fig. 5i). When external recombinant human endostatin was added to wild-type cells, we observed significant increases in histone H3 and its modifications in both PDAC cell lines (Fig. 5j-5k), which were similar to the effect of SPINK1 overexpression. Prior studies suggested several downstream signaling pathways of endostatin [59], including nucleolin as a receptor of endostatin, which is a major nucleolar protein and can transport from the cell surface to the nucleus [60]. After the addition of endostatin, the phosphorylation of nucleolin (NCL) showed acute increases in both PANC-1 and MiaPaCa2 cell lines (Fig. 5l). However, no expression of VEGFR and no changes in the activities of β-Catenin or Src were detected. This finding suggested nucleolin as the mediator of endostatin leading to downstream effects.

Overall, our data propose a novel mechanism by which SPINK1 regulates the epigenomic states in PDAC. We show that SPINK1 interacts with COL18A1, influencing the release of endostatin which subsequently interacts with nucleolin that induces histone alterations.

### The effects of SPINK1 on histone modifications and cancer stemness are also observed in the primary patient samples

To further validate our findings in a clinically relevant setting, we first obtained a few fresh frozen tumor samples collected from PDAC patients recently undergoing surgical resection. We confirmed the presence of tumor cells through H&E staining (Fig. 6a & S6). We then performed co-staining with SPINK1, KRT19 (used as the ductal cell marker) and histone modification marker (H3K4me3, H3K27me3 or H3K27Ac) antibodies on a series of adjacent slides of the samples (Fig. 6b & S6). Supporting the *in vitro* findings, tumor cells with positive SPINK1 staining showed increased levels of histone modification markers (Fig. 6c), while tumor cells with undetectable SPINK1 staining showed limited histone modification marker signal (Fig. 6d), as e.g., shown in the PDAC patient GI1019. Notably, the region with high SPINK1 was composed of a large cell cluster with numerous tumor cells, whereas the region with undetectable SPINK1 was associated with small cell clusters containing few tumor cells. Similar observations were also found in the other three patients (tumor regions with positive SPINK1 staining in Fig. S6a, S6b, and S6c upper panel; tumor region with undetectable SPINK1 staining in Fig. S6c lower panel).

**Figure 6.**
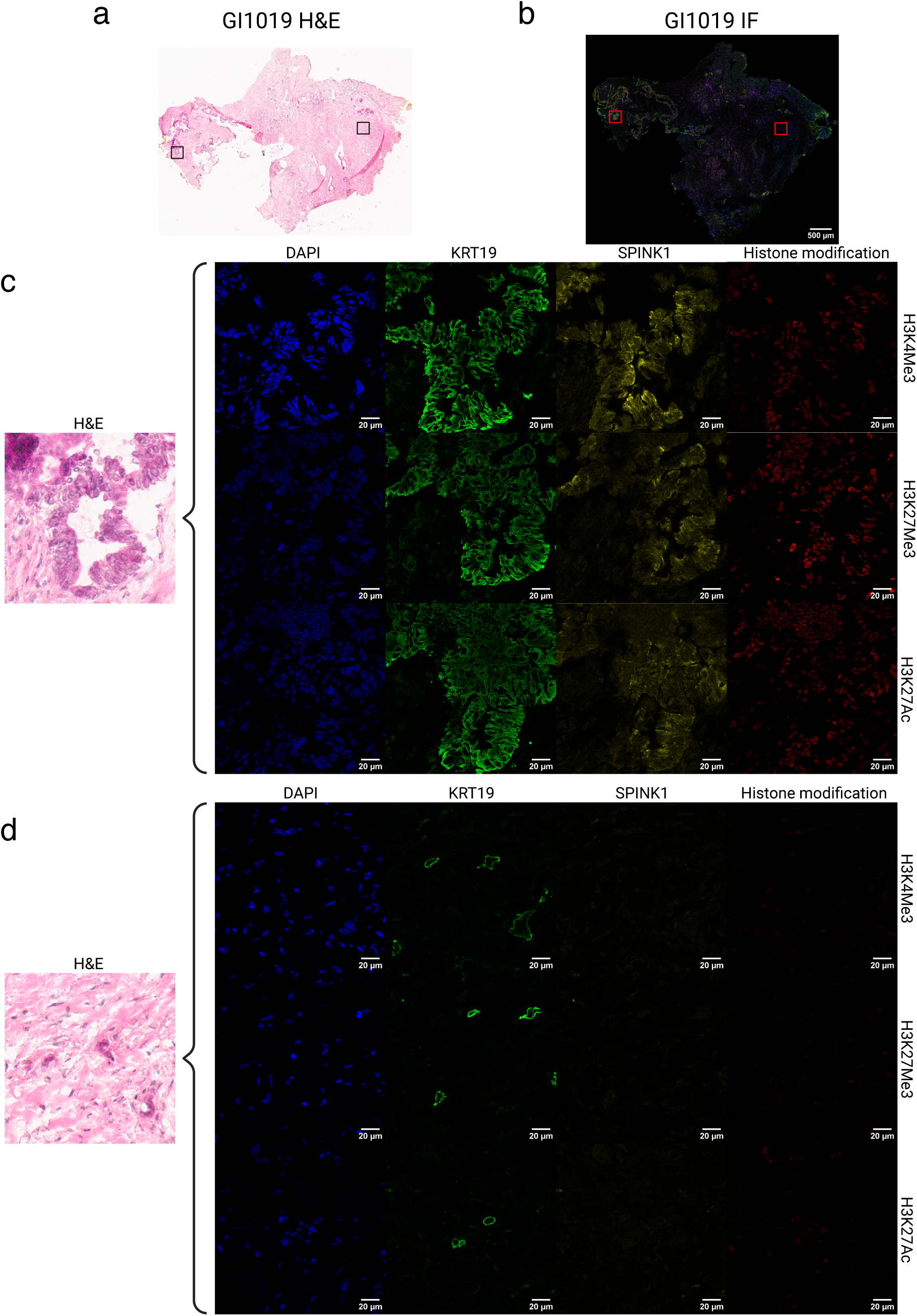
The correlation between SPINK1 and histone modification levels found *in vitro* is also observed in the primary samples of PDAC patients. (a) Overview of H&E staining of the tumor from the patient GI1019 (black boxes marked the selected locations of following representative regions). (b) Overview of immunofluorescence (IF) image of the tumor from the same patient (red boxes marked the selected locations of following representative regions) (c) Representative region with high SPINK1 and KRT19 showed high histone H3 modification markers in the IF image of a PDAC patient. (d) Representative region with low SPINK1 and high KRT19 showed low or no histone H3 modification markers in the IF image of the same PDAC patient.

ALDH1A1, an evolutionarily conserved enzyme, has been extensively studied and commonly accepted as a marker for CSCs across multiple cancers including PDAC [61–64]. Since ALDH1A1 showed a positive correlation to SPINK1 in both the scRNA-seq reference dataset and the STS versus LTS cohort (Fig. 4e-4f), we adopted it as a CSC marker in detecting the relationship between SPINK1 and PDAC cancer stemness in the primary samples. A tissue microarray (TMA) of FFPE PDAC samples was repurchased and co-stained with SPINK1 and ALDH1A1, with KRT19 staining and pathologist-verified H&E used to identify the tumor regions (Fig. 7a & S7). As shown in the four representative PDAC patients in Fig. 7a, tumor cells with completely low SPINK1 levels showed undetectable ALDH1A1 marker (left panel), while tumors with heterogeneous (middle two panels) or high SPINK1 (right panel) showed high ALDH1A1 levels. In the magnified tumor regions selected from the middle two panels, a greater heterogeneity of SPINK1 vs high ALDH1A1 expression (Fig. 7b) agreed with our hypothesized mechanism postulating that endostatin is released into the extracellular environment due to SPINK1-COL18A1 interaction and hence could induce not only autocrine but paracrine downstream effects. Indeed, quantification of all 30 pathologist-verified PDAC sections showed that more than 90% of the cases were explainable based on this hypothesis (Fig. 7c).

**Figure 7.**
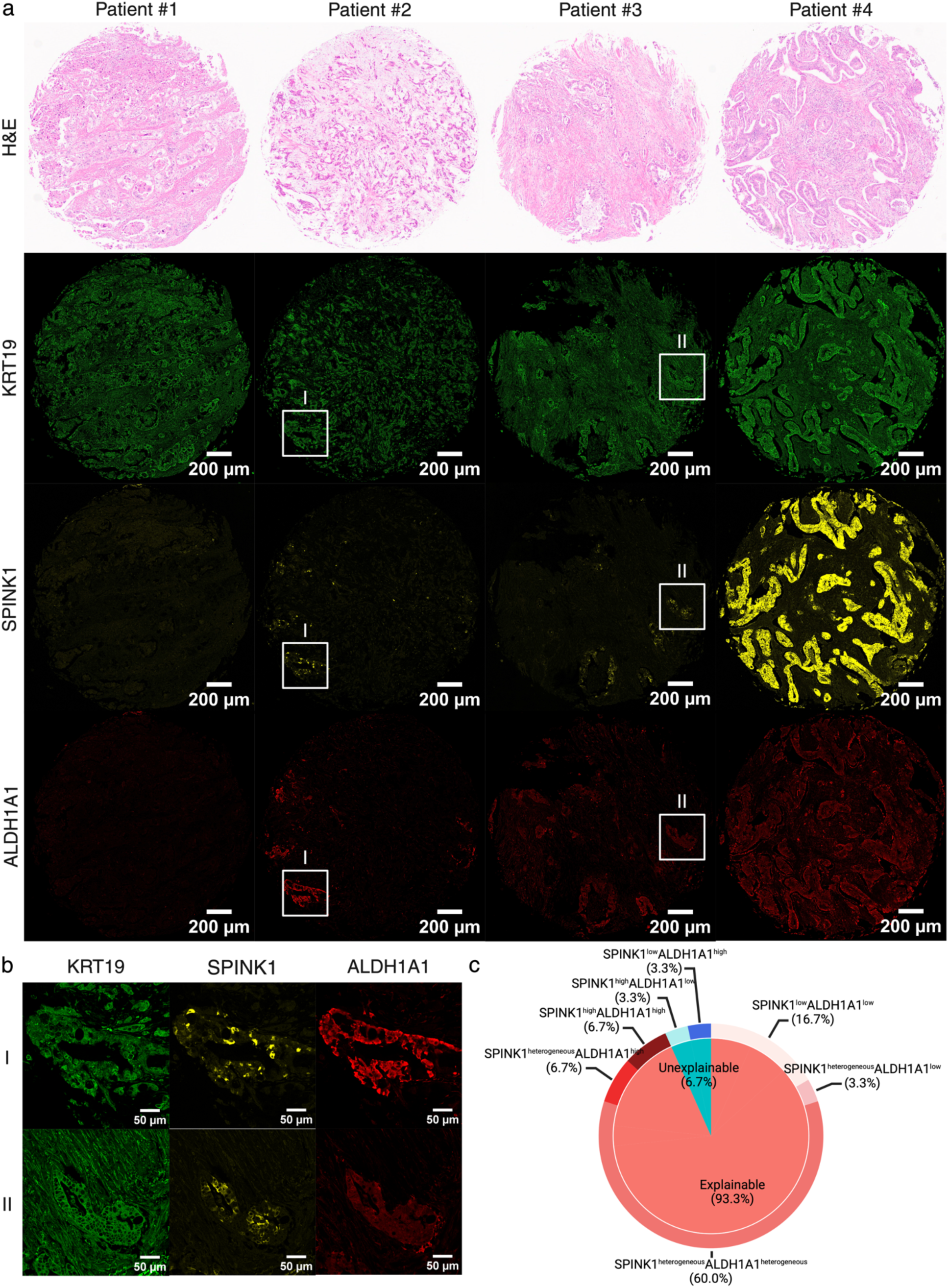
The correlation between SPINK1 and PDAC cancer stemness found *in vitro* is also observed in the primary samples. (a) Representative images showing immunostaining of SPINK1 and stemness marker ALDH1A in TMA. (b) Magnified images from the selected regions (white square windows) from Fig 7a. (c) Pie-donut chart of the correlation between SPINK1 and cancer stemness markers of 30 PDAC patients based on TMA IF staining, which was explainable or unexplainable by our proposed SPINK1 model.

## Discussion

Comparative study between STS and LTS is a common way to identify factors influencing patient outcomes. Previous studies focusing on the analysis of factors determining patient survival, through revealing transcriptomic differences between STS and LTS in PDAC, have been limited by the stringency of defining LTS and STS [65] or hampered by a relatively low number of LTS cases in this very aggressive cancer [66]. TCGA, arguably the largest database focusing on cancer genomics, has only four LTS PDAC cases, with the overall survival (OS) longer than 5 years [67], making it difficult to discover the associated genes. In this study, we overcame this difficulty with a carefully curated patient cohort containing a relatively large number of stringently selected LTS cases (OS ≥ 5 years). By further cross-referencing the bulk transcriptomic result from our cohort with a corrected PDAC scRNA-seq atlas, we found that whereas most genes associated with high expression in STS were primarily expressed in tumor cells (Fig. S1c), two out of five genes linked to long survival were highly expressed in non-tumor cells (Fig. S1d). Notably, one of these two genes, NR4A1, was found to be highly expressed in T-cells, suggesting that the elevated NR4A1 level in T-cells of LTS patients might help prolong their survival, which can be of potential interest for future studies focusing on T-cell-based immunotherapy in PDAC.

The STS-associated gene that was especially highly represented in PDAC patient cells in our single-cell scRNA-seq analysis was SPINK1. Using the trajectory analysis performed on this dataset, we found that the expression of SPINK1 was particularly upregulated in cells emerging during the onset of cancer (the ‘transition’ subcluster) and then remained relatively high during implied cancer progression through the phases characterized by high expression of EMT and cell proliferation associated genes. This result suggested that SPINK1 was potentially involved in enabling diverse cancer related phenotypes, consistent with its previously proposed role as a plasticity related gene in PDAC [68] and other cancers [28, 69]. Cell plasticity is frequently controlled on epigenetic level [70]. We indeed found that genetic perturbation of SPINK1 across different PDAC cell lines strongly influenced H3 chromatin modifications and overall chromatin accessibility in concert with extensive alterations in gene expression. Phenotypically, SPINK1 expression was significantly correlated with the expression of stemness genes both in the PDAC single-cell RNA-seq database and in the micro-array data analysis of our STS versus LTS patient cohort, and the cancer cell colony formation, a key phenotypic feature of cancer stem cells. Furthermore, a correlation between SPINK1 expression and histone modification as well as expression of a key stemness marker was found in PDAC primary tumor sections. Together, these results provided a mechanistic insight into the role of SPINK1 as a plasticity regulator in PDAC and suggested that this role can have a strong negative impact on patient survival.

High PDAC mortality is ascribed to the high invasive and metastatic potential of PDAC tumor cells, enabled by EMT [71]. We thus focused on the possible role of SPINK1 as an STS-associated gene in promoting EMT. SPINK1 has indeed been associated with EMT in several other cancers [72, 73], however, there is limited evidence of a similar relationship between SPINK1 and EMT in PDAC. Our results showed that SPINK1 promoted EMT through modulating the transcription of EMT related genes in PANC-1 cell line, but not in other PDAC cell lines we examined. This suggested that SPINK1 was an indirect regulator of EMT that may enable this phenotype but not directly induce it. This result was consistent with our observations of the association of high SPINK1 expression with cancer cell plasticity and a range of cell phenotypes as discussed above. Our data further suggested that the strong induction of EMT as a specific phenotype might lead to a decrease in SPINK1 expression, possibly reflecting a more differentiated cell state. The additional factors that might affect the adoption of this phenotype in EMT-like state of PANC-1 vs. other cell lines remain to be elucidated. These results suggest that prior studies associating SPINK1 expression with EMT might have reflected a permissive rather than inductive role of this protein.

Our findings prompted us to investigate the putative mechanism of SPINK1-mediated epigenetic control. Although, the canonical function of SPINK1 is that of an inhibitor of trypsin, we found that its role as an epigenetic modifier might depend on a distinct protein domain (Fig. S5h). Prior reports also suggested that SPINK1 may exert its oncogenic effect by activating EGFR and its downstream signaling pathways across several cancers [28, 29, 73–77], due to the structural resemblance with epidermal growth factor (EGF) [78]. In particular, a previous study in PDAC showed the activation of EGFR by rSPINK1 under serum-free condition [56]. However, we found that significant effects of SPINK1 on epigenomic alterations of PDAC cells occurred in complete medium, yet we did not observe that SPINK1 activated EGFR under this condition. These findings suggested the existence of a different, as yet unexplored, mechanism of SPINK1 action to trigger epigenomic changes in PDAC. Using a proximity labeling-based method coupled with mass spectroscopy to screen the candidates, we discovered a new SPINK1 interacting protein COL18A1, and showed that the interaction between SPINK1 and COL18A1 occurred in the Golgi apparatus. We next demonstrated that SPINK1-COL18A1 interaction modulated the cleavage of COL18A1 leading to the release of its active moiety, endostatin. We further showed that endostatin induced the histone changes in PDAC cells, mimicking the effect of perturbations of SPINK1. Moreover, we found that the effect of endostatin was dependent on nucleolin, a protein found in in the nucleolus and other parts of the nucleus and previously shown to promote epigenetic chromatin remodeling [79]. We, therefore, hypothesized a new mechanism of SPINK1-COL18A1-endostatin axis in regulating PDAC epigenomics and cell plasticity, as shown in Fig. 8.

**Figure 8.**
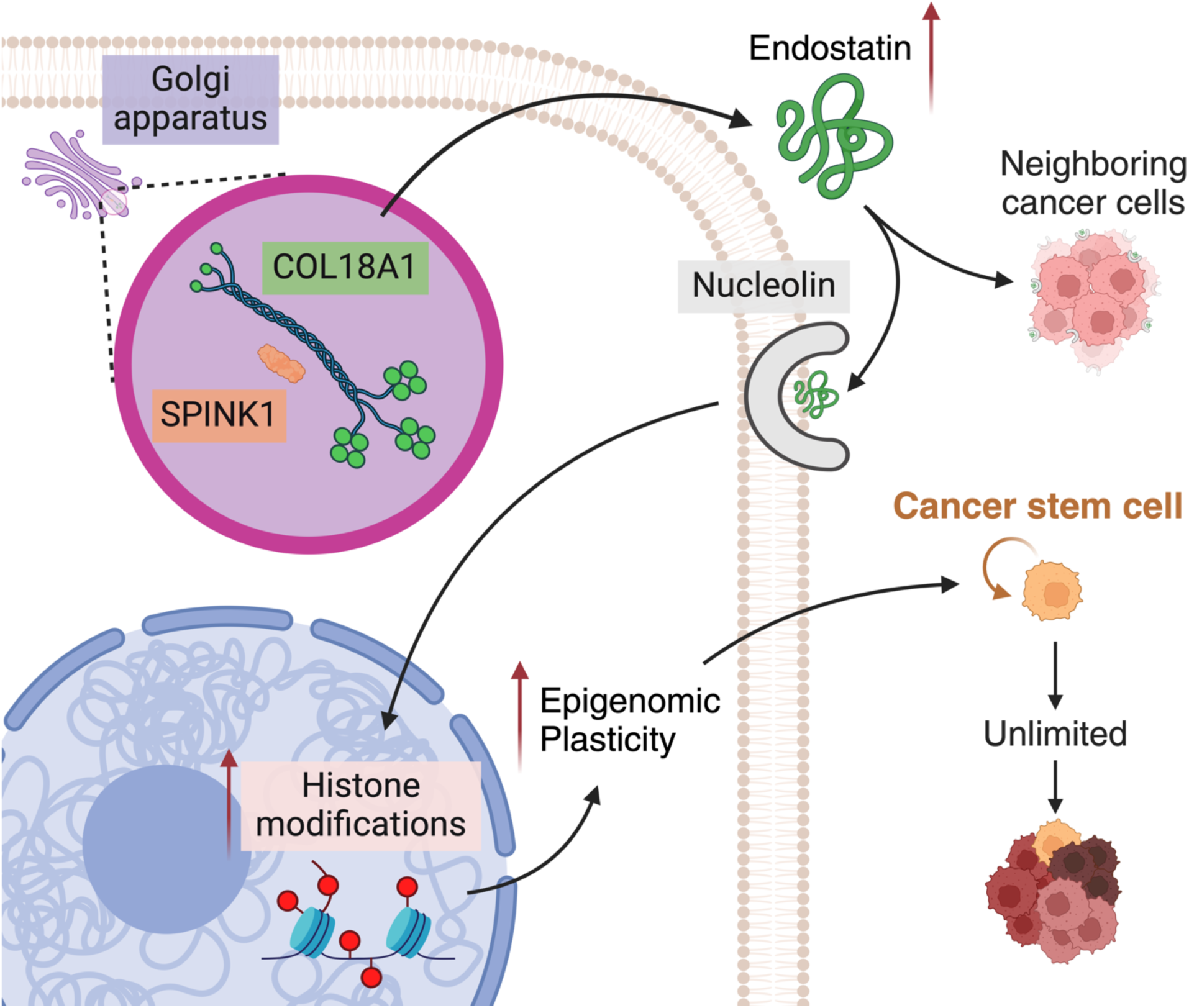
Proposed model of SPINK1 downstream in PDAC. The model shows that SPINK1 increases cancer stemness through interacting with COL18A1 and subsequently changing histone modifications in PDAC. Schematic was created with BioRender.com.

The proposed putative mechanism of SPINK1 action in PDAC may provide further insights relevant to this cancer. In particular, it has been observed that PDAC tumors differ from other solid tumors in terms of having sparse vasculature [80, 81]. And it is reported that PDAC patients with increased tumor vascularity have an improved overall survival, potentially due to a better delivery of anti-cancer chemotherapeutics or immune cells [81, 82]. Endostatin is an angiogenesis inhibitor that specifically reduces endothelial cell proliferation [58]. Based on our model, we hypothesize that an increased endostatin abundance due to the SPINK1-COL18A1 interaction in PDAC cells may be one of the mechanisms contributing to PDAC hypovascularity and poor survival. This effect can be synergistic with SPINK1 and endostatin roles in controlling PDAC cell stemness, making SPINK1 a particularly interesting candidate for therapeutic intervention.

The importance of epigenomic alterations in PDAC progression has been highlighted by multiple studies [8, 9, 83, 84], but the mechanisms regulating these processes remain elusive. Our work supports a critical role of SPINK1 in control of the epigenetic state of tumor cells, the resulting cell plasticity, and thereby patient survival. Although SPINK1 is important in the normal function of the pancreas and is expressed in acinar pancreatic cells, it undergoes strong re-expression as PDAC emerges in ductal-like cancer cells, acquiring a new functional role and operating in a distinct mechanism. The increased plastic state can enable diverse phenotypic outcomes, including EMT, complicate targeting PDAC cells, and thus impact treatment efficacy. Overall, our analysis helps fill the knowledge gap regarding the roles of putative epigenetic modulators in pancreatic cancer progression, addressing an important need for understanding this deadly disease.

## Materials and Methods

### Patient Samples

After approval from institutional review board requirements (IRB 2000022655) at the Department of Surgery, Johns Hopkins University, a registry of PDAC patients undergoing surgery from 1970-2008 was formulated. They were stage-matched and after informed consent, surgical specimens were collected at the Department of Surgery, Johns Hopkins University. All tumors underwent laser microdissection and histological confirmation by expert pathologists (LW and CID). Among these cases, patients with prolonged survival (>5 years; STS) and those with limited survival (<1 year; LTS) were identified. 100 mg of fresh frozen resected specimens were utilized for RNA extraction which was performed through All prep kit (Qiagen, 80204) according to the manufacturer’s protocol. RNA expression levels were detected using Agilent Human Gene Expression 8 × 60 K v2 Microarray Kit.

Another cohort of OCT embedded PDAC tissue samples from 4 PDAC resections at Yale School of Medicine were sectioned, stained with hematoxylin & eosin (H&E) and evaluated by a pancreas pathologist (Dr Marie Roberts) to confirm regions with cancer. All samples were consented and collected under IRB approved protocol (IRB#2000022655). Once confirmed, the samples were processed for immunofluorescence staining. Patient information is given in Supplementary table No 2.

### Cell lines and culture

All human pancreatic cancer cell lines (MiaPaCa2, Capan-1, BxPC-3, PANC-1, HPAF-II) were purchased from the American Type Culture Collection (ATCC) and maintained following the manufacturer’s recommendations.

### Plasmids

hSPINK1 with C terminal ALFA tag in pTwist-Lenti-SFFV-Puro-WPRE backbone (plentiSFFV-hSPINK1-ALFA) was purchased from Twist Bioscience. The control plasmid (plentiSFFV-ALFA) and plasmids with different SPINK1 truncated mutants were constructed by deleting hSPINK1 or the corresponding domain from plentiSFFV-hSPINK1-ALFA with Q5 site-directed mutagenesis kit (NEB, E0554S). For UltraID-based proximity dependent biotinylation, UltraID with N terminal 2×(GGGGS) linker was purchased from Twist Bioscience and inserted into plentiSFFV-hSPINK1-ALFA to generate plentiSFFV-hSPINK1-UltraID and plentiSFFV-hSPINK1SP-UltraID-ALFA using NEBuilder HiFi DNA assembly cloning kit (NEB, E5520S). All constructs were verified by whole plasmid sequencing (Quintara Bioscience). A complete list of all primers used in this study can be found in Supplementary Table 4.

### Tissue microarray

The tissue microarray (TMA) used for immunofluorescence analysis contained both paraffin-embedded normal and pancreatic cancer tissues (TissueArray.Com LLC, MD, USA; Catalog No. PA483-L97) from different individuals. The TMA slide has a total of 48 cores that consist of 38 cases of PDAC, one each of PACC and adenosquamous carcinoma, and 8 normal pancreas tissues, single core per case. After H&E staining and verification by the pathologist, 30 out of 38 cases were confirmed with PDAC tumor present and used together with 8 normal pancreas tissues for the immunofluorescence study. (Fig. S3, where PF-1 to PF-8 represented normal donor tissues and the rest were PDAC tumors).

### Immunofluorescence of tissue microarray

H&E stained and unstained TMA slides containing human PDAC samples and healthy normal controls were obtained from TissueArray.Com (PA483-L97). The H&E-stained TMA slide was inspected by a professional pathologist and samples which could not be confidently classified as PDAC were excluded. The TMA slide for ALDH1A1 staining was obtained as fresh cut and processed within one week upon receiving. For immunofluorescence staining, unstained TMA slides were baked at 60°C for one hour, deparaffinized in xylene, antigen retrieved by heating in citrate buffer, and blocked in 5% normal horse serum. The slide was then incubated with primary antibodies for SPINK1 (H00006690-M01, Abnova) and ALDH1A1 (#64326, Cell Signaling) at 1:100 dilution for 30 min at room temperature. After three rounds of washing with Tris-buffered saline (TBS), the slide was incubated in CF594 labelled anti-mouse (20111-1, Biotium) and AlexaFluor 647 labeled anti-rabbit (A27040, ThermoFisher) secondary antibody at 1:1000 dilution for 30min at room temperature. After three rounds of washing with TBS, the slide was incubated with AlexaFluor 488 labelled KRT-19 antibody (sc-376126 AF488, Santa Cruz Biotechnology) for 30min at room temperature. The slide was then stained with DAPI and washed in PBS for three times and mounted with ProLong Gold antifade mountant (P36934, ThermoFisher).

### CRISPR/Cas9 mediated SPINK1 knockout

Panc-1 and BxPC3 cells were seeded in 24 well plates at a density of 2.5 × 10^4^ cells/well. After attaining 60-70% confluency, the cells were transfected with 500ng of SPINK1 gRNA vector and 500 ng of donor DNA using the non-homology mediated CRISPR knockout kit (Origene, #KN406574) and FuGENE® 4K Transfection Reagent (Promega, #E5911). After 48 hrs of transfection, the cells were split 1:10, and grown additional 3 days; then split 1:10 again twice. The positive selection carried out using puromycin treatment (8ug/ml for Panc-1 cells and 4ug/ml for BxPC3cells) followed by GFP sorting using FACS and subsequently single cell selection was performed. The single-cell selected SPINK1 knockout clones were screened using Western blot analysis. The target sequence for gRNA vector #1: ACTTACCAGATAGACTCAAC and gRNA vector #2: AGGGTCAGCCACATCAATAG. SPINK1 knockout clone was generated from the gRNA vector #2. The scramble construct pCas-Scramble-EF1a-GFP (Origene, #GE100021) was transfected into Panc-1 and BxPC3 cells after reaching 60-70% confluency in a 24 well plate, using and FuGENE® 4K Transfection Reagent (Promega, #E5911). After 48 hrs of transfection, the cells were split 1:10 and grown additional 3 days; then split 1:10 again twice. The positive selection was carried out by GFP sorting using FACS.

### Lentivirus packaging and transduction

All plasmids were packaged into lentivirus by co-transfecting with lentiviral packaging plasmid mix (Cellecta) into HEK293FT cell line (Thermo Fisher Scientific) using lipofectamine 3000 (Thermo Fisher Scientific). The supernatant was 0.22 um filtered and 30x concentrated by lenti-X concentrator (Takara Bio). The concentrated lentivirus was tittered by a qPCR lentivirus titer kit (Applied Biological Materials). The concentrated lentivirus was then added to cells at a MOI of ∼2 in the presence of 10 ug/mL polybrene (Santa Cruz Biotechnology) and incubated for 18∼24 h.

### MTT Assay

The cells were seeded in 96-well plates at a concentration of 2000 cells/100 μl/well and incubated overnight in a CO2 incubator. MTT Assay was carried out at 0, 24, 48 and 72 h time points. After each time point, 10 µl of 5 mg/ml 3-(4,5-dimethyl-2-thiazolyl)-2,5-diphenyl-2H-tetrazolium bromide solution (MTT; Sigma, #475989) was added in each well and incubated at 37°C for 2 h. The MTT formazan crystals were then dissolved with 100 μl/well of DMSO and incubated in dark for 1 hour. Then, the absorbance was measured at 595 nm using iMark™ Microplate Absorbance Reader (BioRad). The cell proliferation due to SPINK1 knockout was determined by normalizing the absorbance reading of 24, 48, and 72 h with 0 h; keeping the absorbance of scramble control cells as 1.

### Immunofluorescence staining of *in vitro* samples

Pancreatic cancer cells were seeded onto a glass-bottom Petri dish (35 mm; Matsunami) at a concentration of 2×10^4^ cells/ml. To enhance cell attachment, all glass-bottom Petri dishes were treated by UV-ozone for 5 minutes before cell seeding. After 3 days of culture, the cells were rinsed briefly in phosphate-buffered saline (PBS) and fixed using 4% paraformaldehyde for 10 min at room temperature. The cells were then washed three times with ice-cold PBS and permeabilized using 0.1% Triton X-100 for 10 min at room temperature. After three times of washing with PBS, the culture dishes were blocked with 3% BSA in PBST (PBS + 0.1% Tween 20) for 1 h at room temperature. The cells were then incubated in 3% BSA in PBST with diluted primary antibodies against COL18A1 (1:200, Atlas Antibodies, Cat# HPA036104) and GOLGA2 (1:200, Proteintech, Cat# 66662-1) at 4 °C overnight. The solution was decanted, and the cells were washed in PBST three times. Subsequently, the cells were incubated with secondary antibodies (1:1000, Thermo Fisher, A32723; 1:1000, Thermo Fisher, A11012 or 1:1000, Jackson ImmuneResearch, 6611-574-215), anti-ALFA Alexa647 antibody (1:500, Nanotag biotechnologies, N1502-AF647-L), and DAPI (1:1000; Sigma, D9542) diluted in 3% BSA in PBST for 1 h at room temperature in the dark. Before imaging, the cells were washed with PBST for three times in the dark. A Stellaris 8 Falcon confocal microscope (Leica, Germany) was used to image the cells, and ImageJ software was adopted to analyze the images.

### Colony Forming Assay

For comparing the colony forming potential between sgCtr versus sgSpink1 and Ctr versus Spink1OE, the cells were seeded at a concentration of 500cells/2ml/well in a 6-well plate and incubated at 37 °C for 2weeks with regular changing of the culture media. After 10-14 days, the colonies formed were fixed with ice-cold methanol for 5 mins followed by staining with 1% crystal violet solution (Sigma-Aldrich, #V5265). The plates were washed with 1X PBS three times and air-dried. Images of the colonies were taken with a scanner (Epson Perfection V39) and the numbers of colonies formed were counted manually.

### Transwell Invasion and Migration Assay

For invasion assay, cells were seeded with serum-free media at a density of 5× 10^4^ into the transwell inserts of 24-well plate Corning BioCoat Matrigel Invasion Chamber (Corning, #354480). For migration assays, 2.5×10^4^ cells were seeded with serum-free media into the 8µm pore transwell inserts of a 24-well plate (Falcon^TM^; #0877121). After 48h of incubation at 37°C, cells that have invaded/ migrated were fixed with 100% methanol for 2 mins and stained with 1% crystal violet solution (Sigma-Aldrich, #V5265). The number of invaded and migrated cells from 3 random fields were counted using a light microscope.

### Western Blot analysis

For protein estimation, Pierce BCA protein Assay Kit (Thermo Fisher Scientific) was used. Protein lysates were prepared using 1x RIPA buffer (Thermo Scientific, Cat# 89900) supplemented with Halt Protease and Phosphatase Inhibitor (Thermo Fisher, Cat# 78442). Then, an equal amount (∼10ug) of each protein sample was denatured with Laemmli sample buffer (Bio-Rad Cat. # 1610747) and 2-mercaptoethanol (Sigma M7522) and boiled at 75°C for 10 min. The denatured samples were resolved in 4–20% Mini-PROTEAN® TGX Stain-Free™ Protein Gels (BioRad, Cat# 4561096) Scientific) and transferred to a PVDF membrane (BioRad, Cat# SLGVR33RS). Membrane blots were blocked with 3% Bovine Serum Albumin (Sigma, Cat# A9647) in 1× tris-buffered saline buffer (dot scientific, Cat# DST60075) supplied with tween (BioRad, Cat# 1706531) (TBST) and probed overnight at 4 °C with the primary antibodies (listed in Supplementary Table 5). Next day, the blots were washed with 1× TBST for 15 min twice and incubated with the appropriate horseradish peroxidase (HRP)-conjugated secondary anti-mouse (GE HealthCare, Cat# NA931) or anti-rabbit antibodies (GE HealthCare, Cat# NA934) for 1h at room temperature. After washing with 1× TBST twice, protein signals were detected with Clarity Western ECL Substrate (BioRad, Cat# 1705061) following the manufacturer’s instructions. For all the immunoblots, GAPDH and β-actin were used as loading controls. The images were all quantified using ImageJ software.

### Treatment by EGF, rSPINK1, endostatin, and TGFβ1

Cells were seeded at 4 × 10^5^ cells/ml in 6-well plates and cultured for 48 h first.

For EGFR signaling test, the control group was changed with fresh medium, while the treatment group was changed with fresh medium added with 10 nM human EGF (PeproTech, Cat# AF-100-15-100UG) or 30 nM recombinant human SPINK1 protein (Biotechne, Cat# 7496-PI-010). After cultured for 10 min or 60 min, cells were washed with ice cold PBS for 1 time and lysed by RIPA buffer supplied with protease and phosphatase inhibitor for subsequent western blot.

As for endostatin function test, the control group was changed with fresh medium, while the treatment group was changed with fresh medium added with 10 μg/ml human endostatin (Genscript, Cat# Z02533). After cultured for 1 day, cells were washed with ice-cold PBS for 1 time and lysed by RIPA buffer supplied with protease and phosphatase inhibitor for subsequent western blot.

As for endostatin signaling test, the control group was changed with fresh medium, while the treatment group was changed with fresh medium added with 10 μg/ml human endostatin. After being cultured for different time periods, cells were washed with ice-cold PBS for 1 time and lysed by RIPA buffer supplied with protease and phosphatase inhibitor for subsequent western blot.

As for inducing EMT test, the control group was changed with fresh medium, while the treatment group was changed with fresh medium added with 10 ng/ml human TGFβ1 recombinant protein (ThermoFisher, Cat# 100-21-2UG). After cultured for 3 days, cells were washed with ice-cold PBS and lysed for western blot, or trypsinized and processed for quantitative RT-PCR.

### Measurement of SPINK1 and endostatin in conditional culture medium

PANC-1 cells were seeded in a 6-well plate for 1 day in complete medium and then cultured in serum-free medium for 8 h. The medium was then collected and centrifuged at 4000xg in Amicon® Ultra-4 Centrifugal Filter Unit (Millipore, Cat# UFC801024) for 15 min, which ended up about 40 times concentration. The concentrated medium was denatured with Laemmli sample buffer (Bio-Rad, Cat. # 1610747) and 2-mercaptoethanol (Sigma, M7522) and underwent western blot to detect SPINK1 and endostatin.

### Measurement of subcellular proteins

Cells were seeded in a 6-well plate and cultured for 2 days. Cells then were lysed and processed following the manufacturer’s protocol of Subcellular Protein Fractionation Kit for Cultured Cells (ThermoFisher, Cat# 78840). The collected proteins were undergone a Western blot analysis.

### RNA isolation and Quantitative RT-PCR

Total RNA was extracted from the cells using the RNeasy minikit (Qiagen, Cat.no. 74104) according to the manufacturer’s protocol. The extracted RNA (15 ng) was subjected to real-time PCR using SYBR™ Green RNA-to-CT™ 1-Step Kit (Applied Biosystems, CA, USA). All samples were normalized to GAPDH. The relative quantification values were calculated using the comparative Ct method (2^−ΔΔCt^). Fold-change expression values were graphically represented using GraphPad Prism 10 (Dotmatics, Boston, MA). Primer sequences are listed in Supplementary Table No 3 (SPINK1and GAPDH primer sequence).

### Bulk RNA sequencing

Total RNA was extracted from SCR and KO cells using RNeasy Mini Kit (Qiagen, Germany, #74104) according to the manufacturer’s protocol. Library preparation and sequencing were performed at Yale Center for Genome Analysis (YCGA), Yale University. The total RNA quality and integrity are determined by nanodrop and Agilent Bioanalyzer gel respectively. mRNA is purified from approximately 200 ng of total RNA with oligo-dT beads and sheared by incubation at 94°C in the presence of Mg (Roche Kapa mRNA Hyper Prep Cat# KR1352). Following first-strand synthesis with random primers, second-strand synthesis, and A-tailing are performed with dUTP for generating strand-specific sequencing libraries. Adapter ligation with 3’ dTMP overhangs are ligated to library insert fragments. Library amplification amplifies fragments carrying the appropriate adapter sequences at both ends. Strands marked with dUTP are not amplified. Indexed libraries are quantified by qRT-PCR using a commercially available kit (Roch KAPA Biosystems Cat# KK4854) and insert size distribution is determined by the Agilent Bioanalyzer. Samples with a yield of ≥ 0.5 ng/ul and a size distribution of 150-300 bp are used for sequencing. *Flow Cell Preparation and Sequencing*: Sample concentrations are normalized to 2 nM and loaded onto an Illumina NovaSeq 6000 flow cell at a concentration that yields 25 million passing filter clusters per sample. Samples are sequenced using 101 bp paired-end sequencing on an Illumina NovaSeq 6000 according to Illumina protocols. The 10bp unique dual index is read during additional sequencing reads that automatically follow the completion of read 1. Data generated during sequencing runs are simultaneously transferred to the YCGA high-performance computing cluster. A positive control (prepared bacteriophage Phi X library) provided by Illumina is spiked into every lane at a concentration of 0.3% to monitor sequencing quality in real-time. Signal intensities are converted to individual base calls during a run using the system’s Real Time Analysis (RTA3) software. Base calls are transferred from the machine’s dedicated personal computer to the Yale High-Performance Computing cluster via a 1 Gigabit network mount for downstream analysis. Primary analysis and sample de-multiplexing were performed using Illumina’s CASAVA 1.8.2 software suite bcl2fastq2/2.20.0-GCCcore-10.2.0. The generated Fastq files were aligned to GRCh38 using Salmon [85]. The resulting quantification files were then analyzed with DEseq2 [86] to identify DEGs (adjusted p value ≤0.05).

### Bulk ATAC sequencing and analysis

Sample processing and sequencing were performed by Active Motif Inc. (Carlsbad, CA, USA). A total of 100,000 cells per sample were cryopreserved in culture media containing FBS and 5% DMSO and sent to Active Motif for ATAC-sequencing. At Active Motif, library preparation was done and sequenced through paired-end42– base pair (bp) sequencing reads format using the Illumina NovaSeq 6000 platform. Adapters were first trimmed with AdpaterRemoval v2 [87]. The trimmed FastQ files were then aligned to GRCh38 using Bowtie2 [88] with the ‘very-sensitive’ setting. After removing PCR duplicates with the MarkDuplicates function from Picard [89][https://broadinstitute.github.io/picard/] and the problematic regions using the ENCODE blacklist [90], peak calling was performed by MACS2 [91]. Regions with differential accessibility were then identified by DiffBind (FDR≤0.05) [92] [http://bioconductor.org/packages/release/bioc/vignettes/DiffBind/inst/doc/DiffBind.pdf].

### Integrated analysis of RNA-seq and ATAC-seq

To analyze ATAC-seq and RNA-seq in an integrative manner, regions with differential accessibility identified from ATAC-seq were annotated to the corresponding genes using the annotatePeak function from ChIPseeker [93]. The genes with differentially accessible regions were then intersected with DEGs identified by DEseq2 from RNA-seq. A gene was considered to have up/downregulated accessibility if all differentially accessible regions annotated to this gene had up/downregulated accessibility. Consensually up/downregulated genes in both RNA-seq and ATAC-seq were then identified and analyzed separately by Enrichr [50] using its website user interface to identify terms enriched from up/downregulated genes.

### Correction of PDAC scRNA-seq reference dataset

The scRNA-seq reference dataset used in this article was curated by Chijimatsu et al. [35] by integrating six published scRNA-seq datasets of PDAC: GSE154778, GSE155698, GSM429355, OUGS, CA001063, and GSE111672.The following issues were identified and corrected from the original PDAC scRNA-seq reference dataset. 1) Log normalized data was mistakenly used as raw counts for CA001063. This issue was corrected by replacing the log normalized data with the corresponding raw counts obtained from CRA001160. 2) Gene names from GSE111672 contains invalid gene names likely causing unintended conversion of the original gene names into date format (e.g. Septin1 to 1-Sept). To identify and correct all affected gene names, the name list containing corrupted gene names was compared to the uncorrupted name list from an unprocessed version of the same dataset from GSM3036909. 3) Genes contained in each dataset does not fully overlap. This problem is likely caused by each dataset being aligned to different versions of the human reference genome (e.g. CD24 is only included in the more recent version of GRCh38) and different criteria being applied to choose genes to be included in the publicly available dataset (e.g. whether to exclude low counts genes or non-protein coding genes). Additionally, the same gene can have different gene names across different datasets due to mixed use of updated and outdated gene symbols (e.g. IL8, the outdated symbol for CXCL8, was used in five out of the six datasets). In the original reference dataset, the union of all genes from the six datasets was included in the final reference dataset and genes not found in certain datasets were assumed to be zero in counts for all cells from the corresponding datasets. For example, cells from datasets aligned to the reference genome not containing CD24 would be assumed to be zero in counts for CD24 even though the true transcription level is unknown. To correct this issue, the intersection rather than the union of the genes was used in the corrected reference dataset. Additionally, for those genes not included in the intersection, the gene symbols were updated, and the intersection was taken again to include those genes excluded due to mixed use of different versions of gene symbols.

### Analysis of PDAC scRNA-seq reference dataset

All analysis in this section was done in Seurat v5 [94] unless otherwise specified. The corrected reference dataset was integrated by RPCA across different datasets using default settings. UMAP [95] was generated by using the first 30 dimensions from the RPCA integrated dimensional reduction with the number of neighbors set to 30. After inspecting corresponding cell type markers, two clusters wrongly labelled in the original reference dataset were corrected (Figure S1). The pseudobulk for each gene in each cell type was then calculated. For further analysis of acinar cells, ductal cells type 1 and ductal cells type 2, the corresponding cells were isolated from the corrected reference dataset and integrated across different patients using Harmony. For the integration by Harmony [96], highly variable genes (HVGs) were first identified for each patient and genes that occurred as HVGs in ≥35 patients were chosen as the final HVGs for the calculation of PCA. Harmony integration was carried out using the first 25 dimensions from the calculated PCA with a random seed set to 5. The neighbor graph was generated by the FindNeighbors function using the first 25 dimensions from the Harmony corrected dimensional reduction and clusters were identified by the FindClusters function with resolution set to 0.3. UMAP was generated using the first 6 dimensions from the Harmony integrated dimensional reduction with the number of neighbors set to 5. Inference of trajectory and pseudotime were carried out in Monocle3 [37–39]. For the inference of trajectory, cells were clustered by the cluster_cells function based on the UMAP from Harmony with the parameter k set to 75 and the trajectory was then inferred by the learn_graph function with close_loop set to FALSE and minimal_branch_len set to 5. The ductal cell type II cluster was further divided into different subclusters: transition (28≤pseudotime≤42), bifurcation (60≤pseudotime≤66.5), high SPINK1 end (cluster 5 from FingClusters), low SPINK1 end (pseudotime≥72). DEG analysis between different subclusters was done by the FindMarkers function from Seurat v5.

### Proximity dependent biotinylation with UltraID

All micro centrifuge tubes used in this section were low-protein binding (Thermo Fisher, 90140). Panc-1 cells overexpressing SPINK1-UltraID-ALFA and SPINK1SP-UltraID-ALFA were allowed to grow to ∼90% confluency in 150 mm Petri dish. On the day of the experiment, cells were incubated in 50 µM biotin for 1h at 37℃. After incubation, cells were washed three times with 10 mL DPBS. 1mL ice-cold lysis buffer (RIPA + 1X Halt^TM^ protease inhibitor cocktail (Thermo Fisher, 78429)) was then added for each 150 mm Petri dish and spread by a cell scraper. The resulting lysate from each plate was then transferred into a micro centrifuge tube and incubated on ice for 30 min with periodic vortexing every 10 min. The lysate was then centrifuged at 13,000xg for 10 min at 4℃ and the supernatant was transferred into a new tube. The protein concentration of each sample was then determined by BCA assay and adjusted to the same. The lysate from each Petri dish was then incubated with 50 µL trypsin-resistant streptavidin beads (MagReSyn, MR-STP002) at 4℃ with gentle shaking overnight. The beads were then pulled down from the lysate and washed at room temperature twice with 1mL lysis buffer, once with 1mL 1M KCl, once with 0.9 mL 0.1M Na2CO3, once with 0.9 mL 2 M urea in 50 mM ammonium bicarbonate (ABC) buffer and twice with 1mL lysis buffer. After the final wash, beads were transferred to a new centrifuge tube with 0.5 mL lysis buffer. Beads were then washed three times with 1 mL 50 mM ABC buffer and resuspended in 200 µL ABC buffer. 10 µL of the beads were aliquoted for quality control. 2 µL 0.5 M TCEP (Thermo Fisher, 77720) was added to the remaining beads in each centrifuge and vortexed to mix. 10 µL freshly prepared 200 mM 2-chloroacetamide (Sigma-Aldrich, C0267-100G) in 50 mM ABC buffer was then added to each centrifuge tube and vortexed to mix. The reaction mixture was shaken in dark at room temperature for 1 h. Beads were then pulled down and washed once with 1 mL 50 mM ABC buffer and resuspended in 200 µL 50 mM ABC buffer containing 250 ng LysC/Trypsin (Promega, V5071). The reaction mixture was then incubated with gentle sharking overnight. Beads were the pulled down and the supernatant was collected and mixed with 25 µL 10% formic acid to quench trypsin digestion. All supernatants were then sent on ice to the Keck proteomics center for analysis by mass spectrometry.

### Co-Immunoprecipitation

For crosslinking co-IP, PANC-1 control and SPINK1-OE cells were seeded in separate 150mm culture dishes (Falcon, Cat# 08-772-6) and cultured until reaching ∼90% of confluency. The medium was discarded, and the cells were washed with ice-cold PBS once. The cells were then added with DSP (Thermo, Cat# 22586; 0.4 mg/ml in PBS, 10 ml per dish) and incubated at 4 °C for 30 min. The crosslinking reaction was stopped by adding 1 ml of PBS containing 0.2 M glycine (Fisher BioReagents, Cat# BP381-1) and incubated at 4 °C for 15 min. After discarding the reaction solution, the cells were gently washed with pre-chilled PBS twice and then lysed using IP Lysis Buffer (Thermo, Cat# 87787; 1 ml per dish) with 1% protease and phosphatase inhibitor cocktail. The cells were scrapped off using a cell lifter and transferred to a 1.5 ml microcentrifuge tube and put on ice for 30 min during which tapping the tube every 5 min to ensure complete lysis. The lysate was next centrifuged at 16,200×g at 4 °C for 10 min. The lysate supernatant was collected and transferred to the equilibrated magnetic ALFA Selector Resin (Nanotag biotechnologies, Cat# N1516) or Selector Control beads (Nanotag biotechnologies, Cat# N0015-S) following the manufacturer’s guide. The lysate-beads mixture was subsequently incubated on a tube rotator at 17 RPM and 4 °C for 1 h. The beads were collected by a magnetic rack and washed using 1×TBS for three times. The bound proteins were eventually eluted from the beads using RIPA buffer supplied with 1% protease and phosphatase inhibitor, Laemmli sample buffer, and 2-mercaptoethanol. The eluted proteins were used for later analysis.

### Immunofluorescence of fresh frozen tissue section

All sections were stored at –80 °C before use and processed within two weeks. For immunofluorescence staining, sections were first warmed up to room temperature for 10min and then fixed in methanol-free 4% PFA (FB002, ThermoFisher) at room temperature for 15 min. Sections were then washed in PBS for three times and permeabilized in PBS with 0.3% Triton-X 100 for 15 min. After permeabilization, sections were washed three times in PBS and blocked in 2% BSA in PBS for 1 hour at room temperature. SPINK1 antibody (1:400, H00006690-M01, Abnova) and antibody for H3K4Me3 (1:200, MA5-11199, ThermoFisher) or H3K27Me3 (1:200, MA5-11198, ThermoFisher) or H3K27Ac (1:200, ab4729, Abcam) modification were then added to the section and incubated for 1 h at room temperature. After three times of washing with PBS, sections were incubated with CF594 labelled anti-mouse antibody (20111-1, Biotium) and AlexaFluor 647 labelled anti-rabbit antibody (A27040, ThermoFisher) at 1:1000 dilution for 1 h at room temperature. The sections were then washed three times with PBS and incubated with AlexaFluor 488 labelled KRT19 antibody (sc-376126 AF488, Santa Cruz Biotechnology) at 1:50 dilution for 1 hour at room temperature. Subsequently, the sections were stained with DAPI and washed three times in PBS and mounted in ProLong Gold mountant (P36934, ThermoFisher). The stained samples were then imaged on a Leica Stellaris 8 Falcon confocal microscope.

### Quantification and statistical analysis

Image quantifications are described here for individual experiments. Student’s t test, chi-square tests, and any other statistical analysis of data were done using GraphPad Prism v.10.0.3 software (GraphPad Software). Pairwise comparisons between groups were made using unpaired Student’s t-tests as indicated. scRNA-seq and patient cohort data were analyzed using R program as indicated above. Scatter or bar plots graphed in GraphPad Prism displayed the standard error mean (SEM) from the mean value. P values less than 0.05 were considered statistically significant and marked with one asterisk mark, meanwhile p values less than 0.01 were marked with two asterisk marks.

## Author’s contribution

H.T, B.S, X.S, A.S, A.L and N.A conceptualized and designed the study; H.T, B.S, X.S, A.S, A.L and N.A developed the methodology; H.T, B.S and X.S conducted the investigations and data analyses. M.R, J.K, E.P, L.W, C.I.D, R.H and C.L.W collected the patient samples and clinical information. P.D performed RT-PCR experiments and data interpretation. A.S, A.C and P.A performed preliminary experiments and data interpretation. R.G.M and M.M performed preliminary bioinformatics analysis. L.H, M.P and O.A.O helped with writing and editing. The original manuscript was drafted by H.T, B.S and X.S, and all authors reviewed and approved the final manuscript. N.A. and A.L acquired funding and supervised the study.

## Acknowledgement

N.A. receives research grant funding from Astex Inc and the Van Andel Research Institute. She is a consultant for and has licensed methylation biomarkers to Cepheid (patent # 10167513). N.A. has served as a consultant to Johnson and Johnson, an advisor to Celgene, and a member of the Scientific Advisory Council to the No Stomach for Cancer Foundation. N.A. also serves as PI on NIH grants 5P30CA016359-42 and 7R01CA185357-05. This work was supported by the NIH grant 7R01CA185357-05, NIH Research Grant P30CA016359 & Yale Cancer Center (YCGA pilot grant). A.L. receives research grant U54 CA209992 from NCI.

We thank the support from Yale Center for Genome Analysis. Sequencing research reported in this publication was supported by the National Institute of General Medical Sciences of the National Institutes of Health under Award Number 1S10OD030363-01A1. We thank the Yale West Campus Imaging Core for the support and assistance in this work. We also thank the MS & Proteomics Resource at Yale University for providing the necessary mass spectrometers and the accompany biotechnology tools funded in part by the Yale School of Medicine and by the Office of The Director, National Institutes of Health (S10OD02365101A1, S10OD019967, and S10OD018034). The funders had no role in study design, data collection and analysis, decision to publish, or preparation of the manuscript.

## References

1. Siegel, R.L., A.N. Giaquinto, and A. Jemal, Cancer statistics, 2024. CA: A Cancer Journal for Clinicians, 2024. 74(1): p. 12–49.

2. Stathis, A. and M.J. Moore, Advanced pancreatic carcinoma: current treatment and future challenges. Nature Reviews Clinical Oncology, 2010. 7(3): p. 163–172.

3. Spiliopoulos, S., et al., Current status of non-surgical treatment of locally advanced pancreatic cancer. World J Gastrointest Oncol, 2021. 13(12): p. 2064–2075.

4. Oh, S.Y., et al., Rare long-term survivors of pancreatic adenocarcinoma without curative resection. World J Gastroenterol, 2015. 21(48): p. 13574–81.

5. Yamamoto, T., et al., Long-term survival after resection of pancreatic cancer: a single-center retrospective analysis. World J Gastroenterol, 2015. 21(1): p. 262–8.

6. Rahib, L., et al., Projecting Cancer Incidence and Deaths to 2030: The Unexpected Burden of Thyroid, Liver, and Pancreas Cancers in the United States. Cancer Research, 2014. 74(11): p. 2913–2921.

7. Yi, J.M., et al., Novel methylation biomarker panel for the early detection of pancreatic cancer. Clin Cancer Res, 2013. 19(23): p. 6544–6555.

8. Lomberk, G., et al., Distinct epigenetic landscapes underlie the pathobiology of pancreatic cancer subtypes. Nature Communications, 2018. 9(1): p. 1978.

9. Lomberk, G., et al., Emerging epigenomic landscapes of pancreatic cancer in the era of precision medicine. Nature Communications, 2019. 10(1): p. 3875.

10. McDonald, O.G., et al., Epigenomic reprogramming during pancreatic cancer progression links anabolic glucose metabolism to distant metastasis. Nature Genetics, 2017. 49(3): p. 367–376.

11. Manoukian, P., et al., Association of epigenetic landscapes with heterogeneity and plasticity in pancreatic cancer. Critical Reviews in Oncology/Hematology, 2025. 206: p. 104573.

12. Sausen, M., et al., Clinical implications of genomic alterations in the tumour and circulation of pancreatic cancer patients. Nature Communications, 2015. 6(1): p. 7686.

13. Waddell, N., et al., Whole genomes redefine the mutational landscape of pancreatic cancer. Nature, 2015. 518(7540): p. 495–501.

14. Bailey, P., et al., Genomic analyses identify molecular subtypes of pancreatic cancer. Nature, 2016. 531(7592): p. 47–52.

15. Wang, S., et al., The molecular biology of pancreatic adenocarcinoma: translational challenges and clinical perspectives. Signal Transduction and Targeted Therapy, 2021. 6(1): p. 249.

16. Ougolkov, A.V., V.N. Bilim, and D.D. Billadeau, Regulation of pancreatic tumor cell proliferation and chemoresistance by the histone methyltransferase enhancer of zeste homologue 2. Clin Cancer Res, 2008. 14(21): p. 6790–6.

17. Ouaïssi, M., et al., High Histone Deacetylase 7 (HDAC7) Expression Is Significantly Associated with Adenocarcinomas of the Pancreas. Annals of Surgical Oncology, 2008. 15(8): p. 2318–2328.

18. Fritsche, P., et al., HDAC2 mediates therapeutic resistance of pancreatic cancer cells via the BH3-only protein NOXA. Gut, 2009. 58(10): p. 1399–409.

19. Kazal, L.A., D.S. Spicer, and R.A. Brahinsky, Isolation of a crystalline trypsin inhibitor-anticoagulant protein from pancreas. J Am Chem Soc, 1948. 70(9): p. 3034–40.

20. Frossard, J.L. and M.A. Morris, Two SPINK1 Mutations Induce Early-Onset Severe Chronic Pancreatitis. Case Rep Gastroenterol, 2017. 11(1): p. 85–88.

21. Abass, M.K., et al., Combined SPINK1 mutations induce early-onset severe chronic pancreatitis in a child with severe obesity. Endocrinol Diabetes Metab Case Rep, 2022. 2022.

22. Muller, N., et al., Natural history of SPINK1 germline mutation related-pancreatitis. EBioMedicine, 2019. 48: p. 581–591.

23. Suzuki, M. and T. Shimizu, Is SPINK1 gene mutation associated with development of pancreatic cancer? New insight from a large retrospective study. EBioMedicine, 2019. 50: p. 5–6.

24. Cazacu, I.M., et al., Pancreatitis-Associated Genes and Pancreatic Cancer Risk: A Systematic Review and Meta-analysis. Pancreas, 2018. 47(9).

25. Piepoli, A., et al., Lack of association between UGT1A7, UGT1A9, ARP, SPINK1 and CFTR gene polymorphisms and pancreatic cancer in Italian patients. World J Gastroenterol, 2006. 12(39): p. 6343–8.

26. Schubert, S., et al., CFTR, SPINK1, PRSS1, and CTRC Mutations Are Not Associated With Pancreatic Cancer in German Patients. Pancreas, 2014. 43(7).

27. Ru, N., et al., SPINK1 mutations and risk of pancreatic cancer in a Chinese cohort. Pancreatology, 2021. 21(5): p. 848–853.

28. Man, K.-F., et al., SPINK1-induced tumor plasticity provides a therapeutic window for chemotherapy in hepatocellular carcinoma. Nature Communications, 2023. 14(1): p. 7863.

29. Chen, F., et al., Targeting SPINK1 in the damaged tumour microenvironment alleviates therapeutic resistance. Nature Communications, 2018. 9(1): p. 4315.

30. Ateeq, B., et al., Therapeutic Targeting of SPINK1-Positive Prostate Cancer. Science Translational Medicine, 2011. 3(72): p. 72ra17–72ra17.

31. Integrated Genomic Characterization of Pancreatic Ductal Adenocarcinoma. Cancer Cell, 2017. 32(2): p. 185–203.e13.

32. Cao, L., et al., Proteogenomic characterization of pancreatic ductal adenocarcinoma. Cell, 2021. 184(19): p. 5031–5052.e26.

33. Biankin, A.V., et al., Expression of S100A2 calcium-binding protein predicts response to pancreatectomy for pancreatic cancer. Gastroenterology, 2009. 137(2): p. 558–68, 568.e1-11.

34. Chen, Q., et al., Dual and triple gene combinations of KRT5, KRT17, and S100A2 identify basal-like subtype of pancreatic ductal adenocarcinoma and correlate with survival outcome. Faseb j, 2024. 38(15): p. e23867.

35. Chijimatsu, R., et al., Establishment of a reference single-cell RNA sequencing dataset for human pancreatic adenocarcinoma. iScience, 2022. 25(8): p. 104659.

36. Peng, J., et al., Single-cell RNA-seq highlights intra-tumoral heterogeneity and malignant progression in pancreatic ductal adenocarcinoma. Cell Research, 2019. 29(9): p. 725–738.

37. Trapnell, C., et al., The dynamics and regulators of cell fate decisions are revealed by pseudotemporal ordering of single cells. Nat Biotechnol, 2014. 32(4): p. 381–386.

38. Qiu, X., et al., Single-cell mRNA quantification and diaerential analysis with Census. Nat Methods, 2017. 14(3): p. 309–315.

39. Qiu, X., et al., Reversed graph embedding resolves complex single-cell trajectories. Nat Methods, 2017. 14(10): p. 979–982.

40. Marstrand-Daucé, L., et al., Acinar-to-Ductal Metaplasia (ADM): On the Road to Pancreatic Intraepithelial Neoplasia (PanIN) and Pancreatic Cancer. Int J Mol Sci, 2023. 24(12).

41. Zhang, J., et al., Role of the translationally controlled tumor protein in DNA damage sensing and repair. Proc Natl Acad Sci U S A, 2012. 109(16): p. E926–33.

42. Boucher, D., et al., hSSB2 (NABP1) is required for the recruitment of RPA during the cellular response to DNA UV damage. Sci Rep, 2021. 11(1): p. 20256.

43. Niehrs, C. and A. Schäfer, Active DNA demethylation by Gadd45 and DNA repair. Trends Cell Biol, 2012. 22(4): p. 220–7.

44. Yang, Y., et al., Transcription Factor C/EBP Homologous Protein in Health and Diseases. Front Immunol, 2017. 8: p. 1612.

45. Diaferia, G.R., et al., Dissection of transcriptional and cis-regulatory control of diaerentiation in human pancreatic cancer. Embo j, 2016. 35(6): p. 595–617.

46. Loh, C.Y., et al., The E-Cadherin and N-Cadherin Switch in Epithelial-to-Mesenchymal Transition: Signaling, Therapeutic Implications, and Challenges. Cells, 2019. 8(10).

47. Usman, S., et al., Vimentin Is at the Heart of Epithelial Mesenchymal Transition (EMT) Mediated Metastasis. Cancers (Basel), 2021. 13(19).

48. Zhang, P., Y. Sun, and L. Ma, ZEB1: at the crossroads of epithelial-mesenchymal transition, metastasis and therapy resistance. Cell Cycle, 2015. 14(4): p. 481–7.

49. Ghandi, M., et al., Next-generation characterization of the Cancer Cell Line Encyclopedia. Nature, 2019. 569(7757): p. 503-508.

50. Kuleshov, M.V., et al., Enrichr: a comprehensive gene set enrichment analysis web server 2016 update. Nucleic Acids Res, 2016. 44(W1): p. W90–7.

51. Barski, A., et al., High-resolution profiling of histone methylations in the human genome. Cell, 2007. 129(4): p. 823–37.

52. Chen, K., et al., Broad H3K4me3 is associated with increased transcription elongation and enhancer activity at tumor-suppressor genes. Nat Genet, 2015. 47(10): p. 1149–57.

53. Creyghton, M.P., et al., Histone H3K27ac separates active from poised enhancers and predicts developmental state. Proc Natl Acad Sci U S A, 2010. 107(50): p. 21931–6.

54. Batlle, E. and H. Clevers, Cancer stem cells revisited. Nature Medicine, 2017. 23(10): p. 1124–1134.

55. Loh, J.-J. and S. Ma, Hallmarks of cancer stemness. Cell Stem Cell, 2024. 31(5): p. 617–639.

56. Ozaki, N., et al., Serine Protease Inhibitor Kazal Type 1 Promotes Proliferation of Pancreatic Cancer Cells through the Epidermal Growth Factor Receptor. Molecular Cancer Research, 2009. 7(9): p. 1572–1581.

57. Kubitz, L., et al., Engineering of ultraID, a compact and hyperactive enzyme for proximity-dependent biotinylation in living cells. Communications Biology, 2022. 5(1): p. 657.

58. O’Reilly, M.S., et al., Endostatin: An Endogenous Inhibitor of Angiogenesis and Tumor Growth. Cell, 1997. 88(2): p. 277–285.

59. Poluzzi, C., R.V. Iozzo, and L. Schaefer, Endostatin and endorepellin: A common route of action for similar angiostatic cancer avengers. Adv Drug Deliv Rev, 2016. 97: p. 156–73.

60. Shi, H., et al., Nucleolin is a receptor that mediates antiangiogenic and antitumor activity of endostatin. Blood, 2007. 110(8): p. 2899–2906.

61. Bo, Y., et al., Leveraging intracellular ALDH1A1 activity for selective cancer stem-like cell labeling and targeted treatment via in vivo click reaction. Proceedings of the National Academy of Sciences, 2023. 120(36): p. e2302342120.

62. Mizukami, T., et al., Immunohistochemical analysis of cancer stem cell markers in pancreatic adenocarcinoma patients after neoadjuvant chemoradiotherapy. BMC Cancer, 2014. 14(1): p. 687.

63. Wang, J., et al., Phosphorylation-dependent regulation of ALDH1A1 by Aurora kinase A: insights on their synergistic relationship in pancreatic cancer. BMC Biology, 2017. 15(1): p. 10.

64. Sergeant, G., et al., Role of cancer stem cells in pancreatic ductal adenocarcinoma. Nature Reviews Clinical Oncology, 2009. 6(10): p. 580–586.

65. Bhardwaj, A., et al., Deeper insights into long-term survival heterogeneity of pancreatic ductal adenocarcinoma (PDAC) patients using integrative individual- and group-level transcriptome network analyses. Scientific Reports, 2022. 12(1): p. 11027.

66. Chen, R., et al., Stromal galectin-1 expression is associated with long-term survival in resectable pancreatic ductal adenocarcinoma. Cancer Biology & Therapy, 2012. 13(10): p. 899–907.

67. Katsuta, E., et al., A prognostic score based on long-term survivor unique transcriptomic signatures predicts patient survival in pancreatic ductal adenocarcinoma. Am J Cancer Res, 2021. 11(9): p. 4294–4307.

68. Ji, Y., Q. Xu, and W. Wang, Single-cell transcriptome reveals the heterogeneity of malignant ductal cells and the prognostic value of REG4 and SPINK1 in primary pancreatic ductal adenocarcinoma. PeerJ, 2024. 12: p. e17350.

69. Tiwari, R., et al., Androgen deprivation upregulates SPINK1 expression and potentiates cellular plasticity in prostate cancer. Nature Communications, 2020. 11(1): p. 384.

70. Feinberg, A.P. and A. Levchenko, Epigenetics as a mediator of plasticity in cancer. Science, 2023. 379(6632): p. eaaw3835.

71. Yang, J., Y. Liu, and S. Liu, The role of epithelial-mesenchymal transition and autophagy in pancreatic ductal adenocarcinoma invasion. Cell Death Dis, 2023. 14(8): p. 506.

72. Ying, H.Y., et al., Serine protease inhibitor Kazal type 1 (SPINK1) downregulates E-cadherin and induces EMT of hepatoma cells to promote hepatocellular carcinoma metastasis via the MEK/ERK signaling pathway. J Dig Dis, 2017. 18(6): p. 349–358.

73. Wang, C., et al., Serine protease inhibitor Kazal type 1 promotes epithelial-mesenchymal transition through EGFR signaling pathway in prostate cancer. Prostate, 2014. 74(7): p. 689–701.

74. Ateeq, B., et al., Therapeutic targeting of SPINK1-positive prostate cancer. Sci Transl Med, 2011. 3(72): p. 72ra17.

75. Mehner, C., et al., Serine protease inhibitor Kazal type 1 (SPINK1) drives proliferation and anoikis resistance in a subset of ovarian cancers. Oncotarget; Vol 6, No 34, 2015.

76. Marchbank, T., A. Mahmood, and R.J. Playford, Pancreatic secretory trypsin inhibitor causes autocrine-mediated migration and invasion in bladder cancer and phosphorylates the EGF receptor, Akt2 and Akt3, and ERK1 and ERK2. Am J Physiol Renal Physiol, 2013. 305(3): p. F382–9.

77. Ying, H.Y., et al., *Serine protease inhibitor Kazal type 1 (SPINK1) downregulates E-*cadherin and induces EMT of hepatoma cells to promote hepatocellular carcinoma metastasis via the MEK/ERK signaling pathway. Journal of Digestive Diseases, 2017. 18(6): p. 349–358.

78. Hunt, L.T., W.C. Barker, and M.O. Dayhom, Epidermal growth factor: Internal duplication and probable relationship to pancreatic secretory trypsin inhibitor. Biochemical and Biophysical Research Communications, 1974. 60(3): p. 1020–1028.

79. Angelov, D., et al., Nucleolin is a histone chaperone with FACT-like activity and assists remodeling of nucleosomes. Embo j, 2006. 25(8): p. 1669–79.

80. Olive, K.P., et al., Inhibition of Hedgehog Signaling Enhances Delivery of Chemotherapy in a Mouse Model of Pancreatic Cancer. Science, 2009. 324(5933): p. 1457–1461.

81. Provenzano, Paolo P., et al., Enzymatic Targeting of the Stroma Ablates Physical Barriers to Treatment of Pancreatic Ductal Adenocarcinoma. Cancer Cell, 2012. 21(3): p. 418–429.

82. Katsuta, E., et al., Pancreatic adenocarcinomas with mature blood vessels have better overall survival. Scientific Reports, 2019. 9(1): p. 1310.

83. McDonald, O.G., et al., Epigenomic reprogramming during pancreatic cancer progression links anabolic glucose metabolism to distant metastasis. Nat Genet, 2017. 49(3): p. 367–376.

84. Pandey, S., V.K. Gupta, and S.P. Lavania, Role of epigenetics in pancreatic ductal adenocarcinoma. Epigenomics, 2023. 15(2): p. 89–110.

85. Patro, R., et al., Salmon provides fast and bias-aware quantification of transcript expression. Nat Methods, 2017. 14(4): p. 417–419.

86. Love, M.I., W. Huber, and S. Anders, Moderated estimation of fold change and dispersion for RNA-seq data with DESeq2. Genome Biol, 2014. 15(12): p. 550.

87. Schubert, M., S. Lindgreen, and L. Orlando, AdapterRemoval v2: rapid adapter trimming, identification, and read merging. BMC Res Notes, 2016. 9: p. 88.

88. Langmead, B. and S.L. Salzberg, Fast gapped-read alignment with Bowtie 2. Nat Methods, 2012. 9(4): p. 357–9.

89. Institute, B., Picard Toolkit. 2019: GitHub Repository.

90. Amemiya, H.M., A. Kundaje, and A.P. Boyle, The ENCODE Blacklist: Identification of Problematic Regions of the Genome. Sci Rep, 2019. 9(1): p. 9354.

91. Zhang, Y., et al., Model-based analysis of ChIP-Seq (MACS). Genome Biol, 2008. 9(9): p. R137.

92. Stark, R.B., Gord, DiaBind: Diaerential binding analysis of ChIP-Seq peak data. 2011: Bioconductor.

93. Wang, Q., et al., Exploring Epigenomic Datasets by ChIPseeker. Curr Protoc, 2022. 2(10): p. e585.

94. Hao, Y., et al., Dictionary learning for integrative, multimodal and scalable single-cell analysis. Nat Biotechnol, 2024. 42(2): p. 293–304.

95. McInnes, L.H., John; Saul, Nathaniel; Großberger, Lukas, UMAP: Uniform Manifold Approximation and Projection. Journal of Open Source Software, 2018. 3(29): p. 861.

96. Korsunsky, I., et al., Fast, sensitive and accurate integration of single-cell data with Harmony. Nat Methods, 2019. 16(12): p. 1289–1296.

