## Supplemental Figures & Tables for "SPINK1-COL18A1 Crosstalk Shapes Epigenome and Drives Cancer Stemness in Pancreatic Ductal Adenocarcinoma"

### Supplementary Data

Supplementary Table 1. Clinical information of STS-LTS patient cohort. Data of average survival and average age are presented as the mean ± s.e.m. and analyzed using a two-tailed unpaired Student's *t*-test. Data of gender, race, and location are presented as the count number (percentage in the group) and analyzed using Chi-square test for contingency table. Data of grade is presented as the count number (percentage in the group) and analyzed using Mann Whitney (Wilcoxon) test for unpaired data.

| | STS (n = 25) | LTS (n = 19) | P-value |
|---|---|---|---|
| **Average Survival** | 0.38 ± 0.02 Years | 8.93 ± 0.97 Years | < 0.01 |
| **Average age** | 64.6 ± 2.0 Years | 63.3 ± 2.2 Years | 0.68 |
| **Gender** | | | |
| **Female** | 11 (44%) | 10 (53%) | 0.57 |
| **Male** | 14 (56%) | 9 (47%) | |
| **Race** | | | |
| **White** | 21 (84%) | 17 (89%) | 0.66 |
| **Black** | 3 (12%) | 2 (11%) | |
| **Others** | 1 (4%) | 0 (0%) | |
| **Grade** | | | |
| **1** | 0 (0%) | 2 (11%) | 0.02 |
| **2** | 10 (40%) | 11 (58%) | |
| **3** | 14 (56%) | 5 (26%) | |
| **4** | 1 (4%) | 0 (0%) | |
| **Unknown** | 0 (0%) | 1 (5%) | |
| **Location** | | | |
| **Head** | 17 (68%) | 17 (89.5%) | 0.20 |
| **Body** | 0 (0%) | 1 (5%) | |
| **Tail** | 5 (20%) | 0 (0%) | |
| **Overlapping lesion** | 1 (4%) | 1 (5%) | |
| **Body, Tail** | 1 (4%) | 0 (0%) | |
| **Head, Body, Tail** | 1 (4%) | 0 (0%) | |

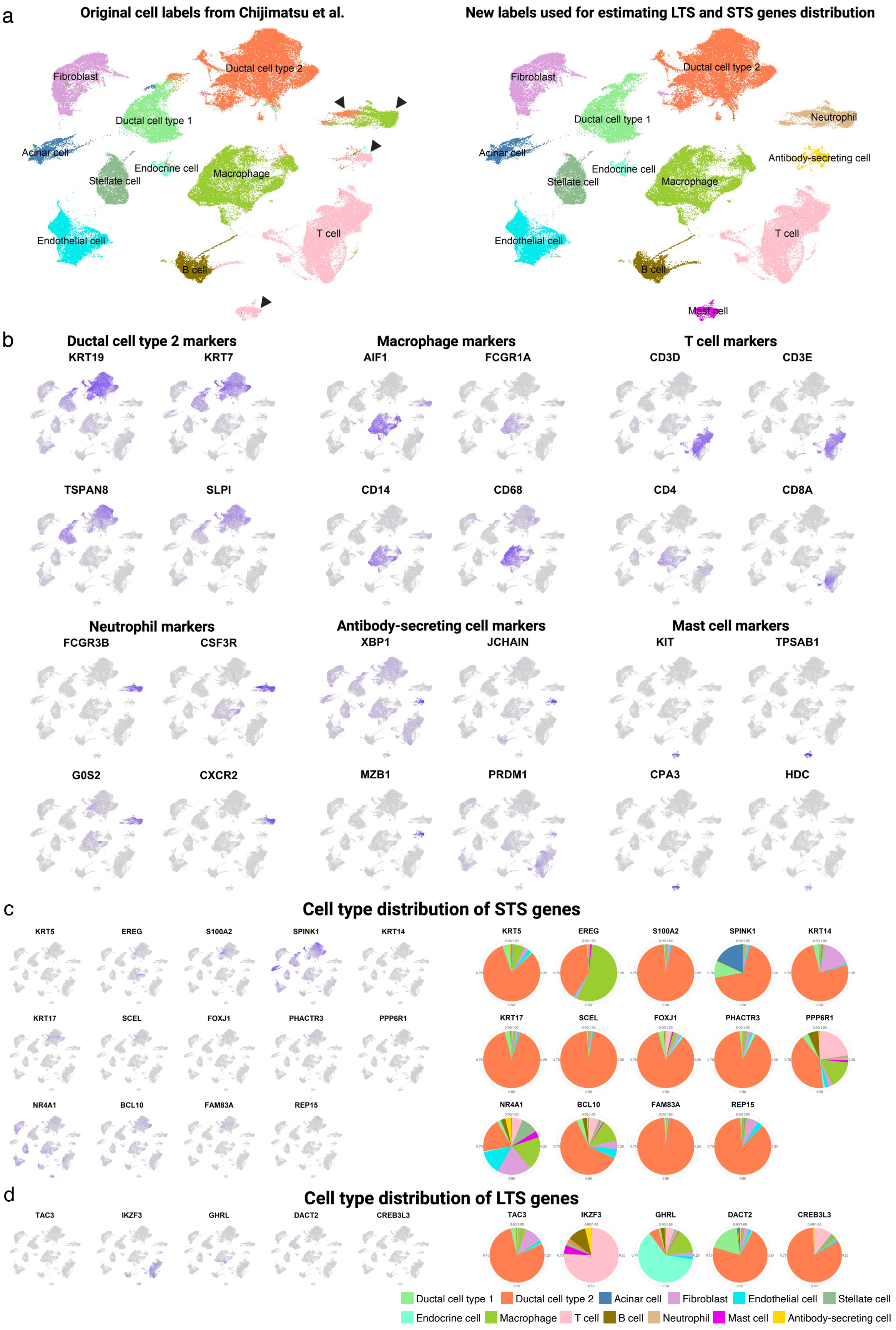
a
Original cell labels from Chijimatsu et al.
Fibroblast
Ductal cell type 2
Ductal cell type 1
Acinar cell
Endocrine cell
Stellate cell
Macrophage
Endothelial cell
T cell
B cell
New labels used for estimating LTS and STS genes distribution
Fibroblast
Ductal cell type 2
Ductal cell type 1
Neutrophil
Acinar cell
Endocrine cell
Antibody-secreting cell
Stellate cell
Macrophage
Endothelial cell
T cell
B cell
Mast cell
b
Ductal cell type 2 markers
KRT19
KRT7
TSPAN8
SLPI
Macrophage markers
AIF1
FCGR1A
CD14
CD68
T cell markers
CD3D
CD3E
CD4
CD8A
Neutrophil markers
FCGR3B
CSF3R
G0S2
CXCR2
Antibody-secreting cell markers
XBP1
JCHAIN
MZB1
PRDM1
Mast cell markers
KIT
TPSAB1
CPA3
HDC
c
Cell type distribution of STS genes
KRT5
EREG
S100A2
SPINK1
KRT14
KRT17
SCEL
FOXJ1
PHACTR3
PPP6R1
NR4A1
BCL10
FAM83A
REP15
d
Cell type distribution of LTS genes
TAC3
IKZF3
GHRL
DACT2
CREB3L3
Ductal cell type 1
Ductal cell type 2
Acinar cell
Fibroblast
Endothelial cell
Stellate cell
Endocrine cell
Macrophage
T cell
B cell
Neutrophil
Mast cell
Antibody-secreting cell

Supplementary Figure 1. Re-cluster of scRNA seq data from the reference database and the projections of the shortlisted genes identified from STS_LTS patient cohort. a) Comparison of new labels of clusters in this study to the original annotations from the work of Chijimatsu et al. b) UMAP of cell markers used to determine cell types of the clusters, leading to the difference of annotations compared to the original study. c) UMAP projections of candidate genes upregulated in STS and their cell type distribution. d) UMAP projections of candidate genes upregulated in LTS and their cell type distribution.

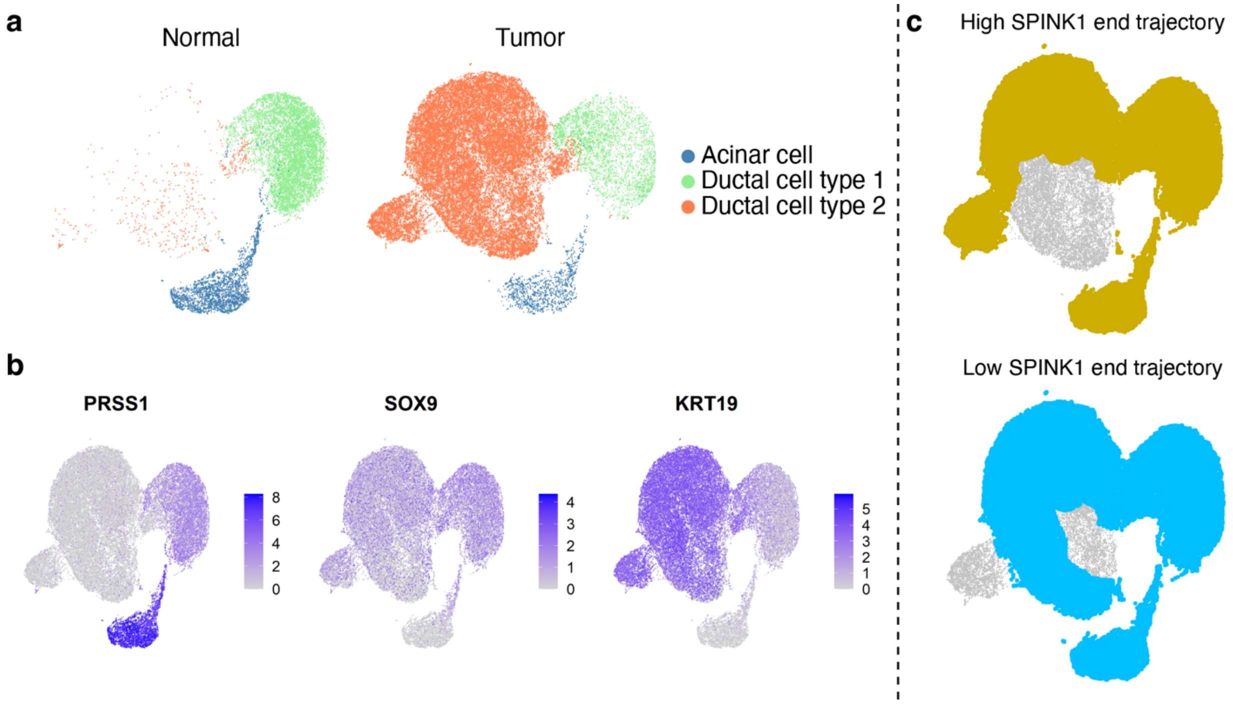


Supplementary Figure 2. The identification of the tumor cell cluster and the trajectory-covered areas. a) Clustering based on cell types. b) UMAP of marker genes used for cell type classification. c) The coverage of cells used for tracking SPINK1 levels along the two trajectories (upper panel for high SPINK1 end trajectory; lower panel for low SPINK1 end trajectory).

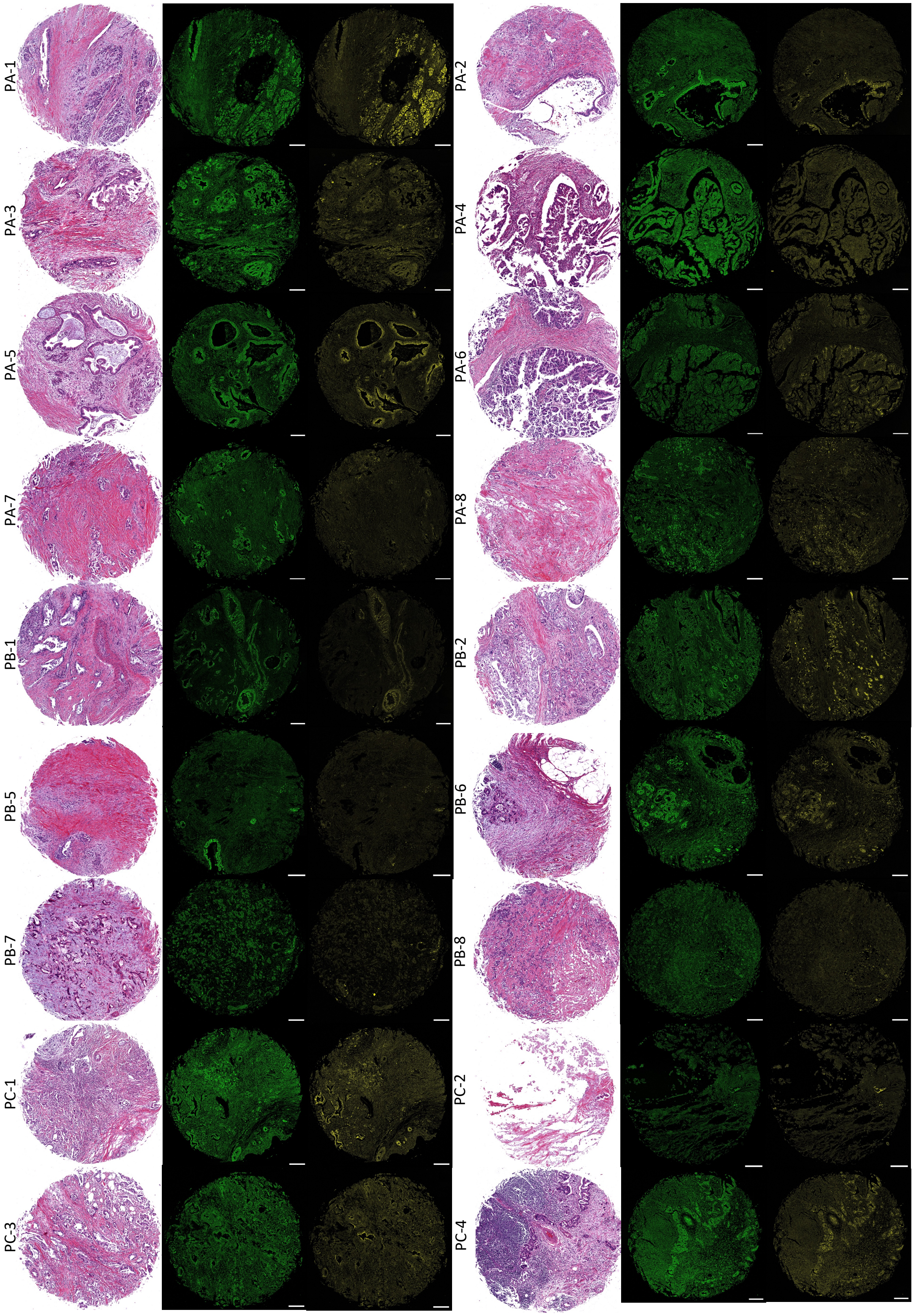
PA-1
PA-2
PA-3
PA-4
PA-5
PA-6
PA-7
PA-8
PB-1
PB-2
PB-5
PB-6
PB-7
PB-8
PC-1
PC-2
PC-3
PC-4

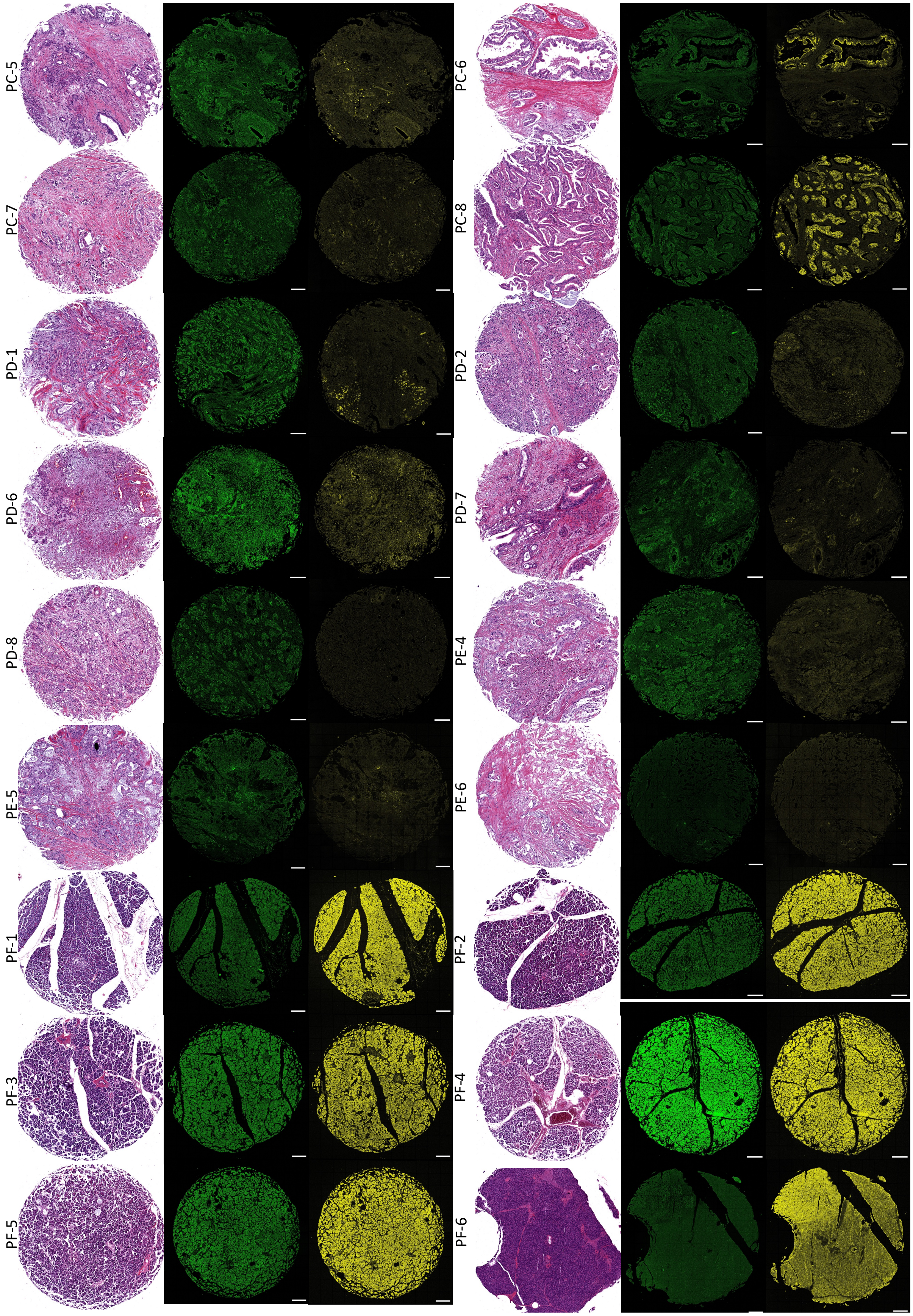
PC-5
PC-6
PC-7
PC-8
PD-1
PD-2
PD-6
PD-7
PD-8
PE-4
PE-5
PE-6
PF-1
PF-2
PF-3
PF-4
PF-5
PF-6

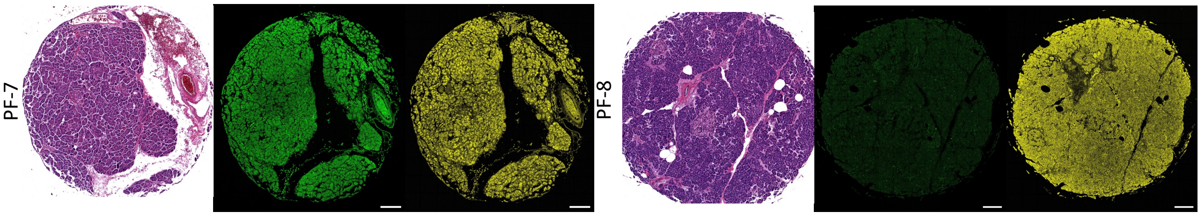


Supplementary Figure 3. H&E and immunofluorescence staining of KRT19 (green) and SPINK1 (yellow) for all PDAC patient samples in TMA. The scale bar is 200μm.

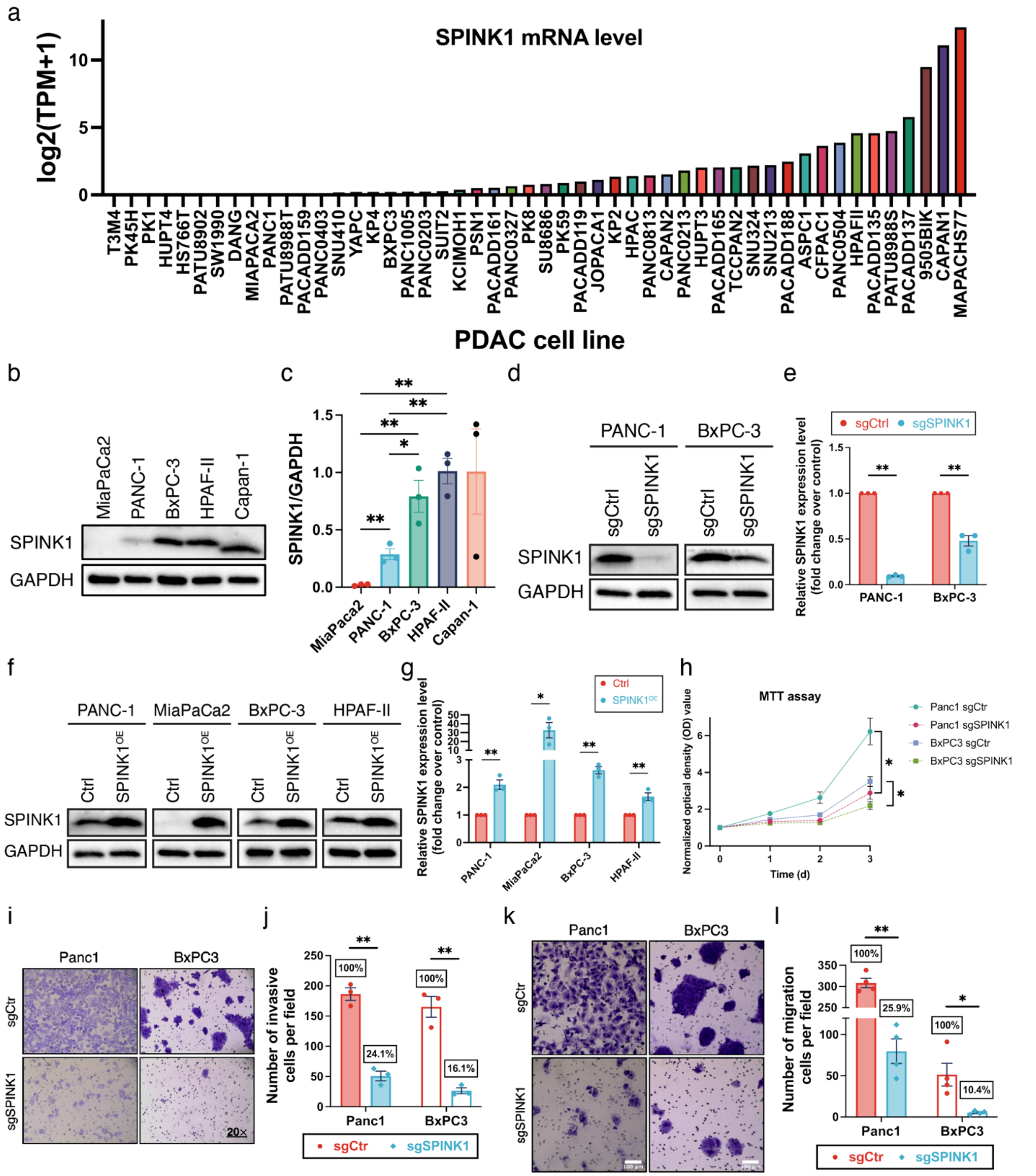


Supplementary Figure 4. The baseline SPINK1 levels and SPINK1 perturbations in PDAC cell lines and the functional assays showing the effect of SPINK1 knock-out. (a) Expression TPM values of SPINK1 in PDAC cell lines according to DepMap Public 24Q2 database. (b) Western blot showing the baseline level of SPINK1 in PDAC cell lines (quantification in c). (d) Western blot showing the SPINK1 levels in PANC-1 and BxPC3 cell lines after CRISPR knockout (quantification in e). (f) Western blot showing the SPINK1 levels in PDAC cell lines after lentiviral overexpression of SPINK1 (quantification in g). (h) MTT assay showing the relative cell viability of SPINK1-knockout cells for proliferation rate analysis. (i) Transwell assay showing the migration ability after SPINK1 knock out. (j) Quantification of transwell migration assay. (k) Invasion assay showing the invasion ability after SPINK1 knock out. (l) Quantification of invasion assay.

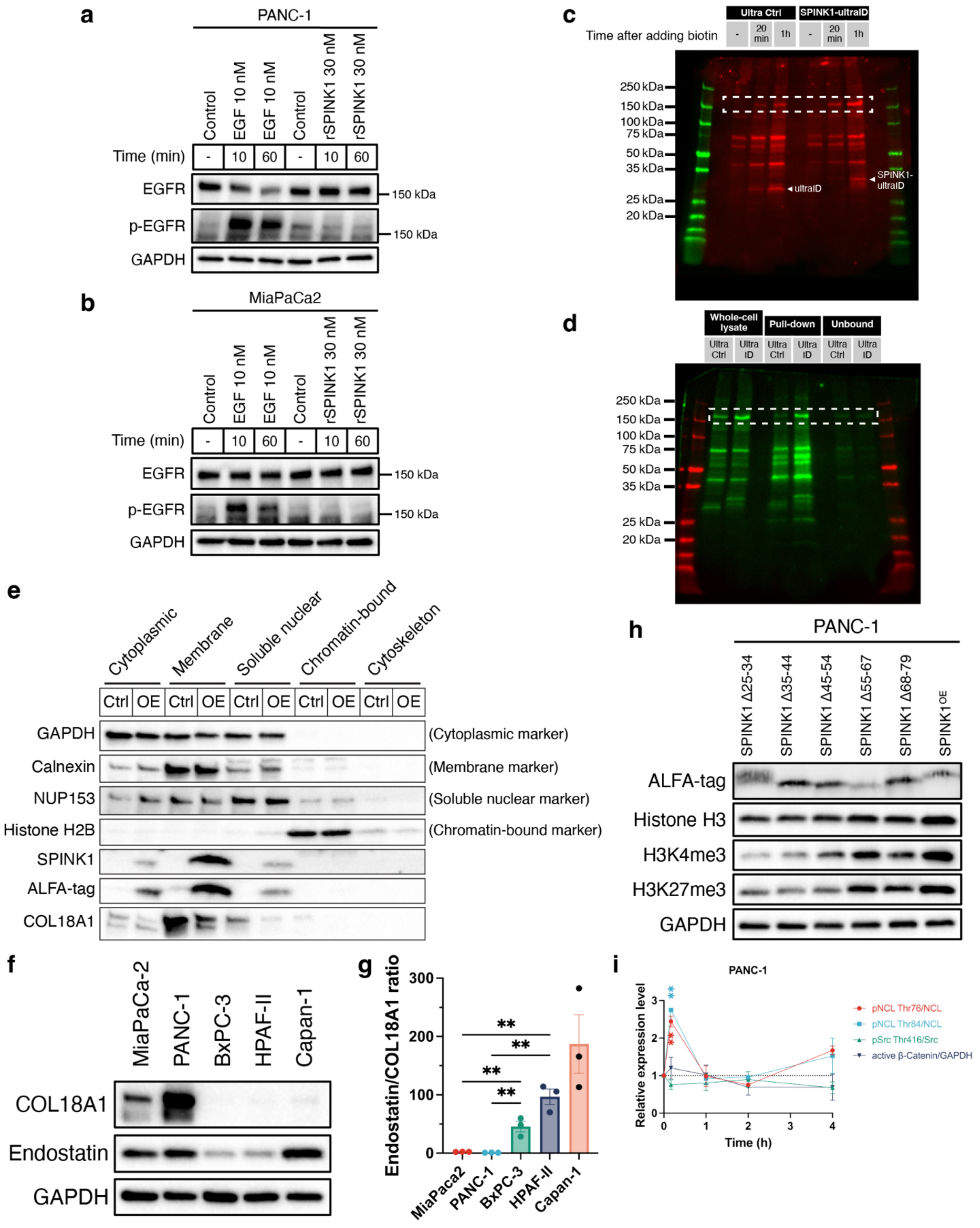

Supplementary Figure 5. SPINK1 showed no significant effect on EGFR but interaction with COL18A1 in PDAC cells. a) Western blot showing the changes in the phosphorylation of EGFR after the treatment with EGF or rSPINK1 in PANC-1 wildtype cells. b) Western blot showing the changes in the phosphorylation of EGFR after the treatment with EGF or rSPINK1 in MiaPaCa2 wildtype cells. c) Western result showing the biotinylated protein bands from ultra Ctrl and SPINK1-ultaID cells after adding biotin with the mark ladder. d) Western result with the ladder showing the biotinylated proteins before and after pull-down using streptavidin beads and the pull-downed proteins. e) Western blot showing the distributions of SPINK1 and COL18A1 in different cellular compartments using PANC-1 Ctrl and SPINK1-OE cells. f) Western blot showing the baseline levels of COL18A1 and endostatin in five examined PDAC cell lines (quantification in g). h) Western blot showing the effect of expression of various partially deleted SPINK1 on histone modifications compared to overexpression of complete SPINK1. i) Quantification for Fig. 5l.

Supplementary Table 2. Patient information for fresh frozen PDAC resection tissue utilized for IF staining of SPINK1 and histone modifications

| PTID# GI | Cancer Location | Histology | Stage | Grade | Tumor Size (cm) | Age | Gender | Comment from Pathologist |
|---|---|---|---|---|---|---|---|---|
| 1019 | Pancreas | Adenocarcinoma | IIB | Invasive Mod. Diff | 4.2 | 80 | F | Confirmed Adenocarcinoma |
| 906 | Pancreas | Adenocarcinoma | III | Invasive Mod. To Poorly Diff. | 4.3 | 71 | M | Confirmed Adenocarcinoma |
| 1064 | Pancreas | Adenocarcinoma | III | Invasive Mod. Diff | 4.5 | 54 | F | Confirmed Adenocarcinoma |
| 796 | Pancreas | Adenocarcinoma | IIB | Invasive Mod. Diff. w/ Mucin | 7.2 | 72 | M | Confirmed Adenocarcinoma |

Supplementary Table 3. Primer sequences used for qPCR:

| Genes | Primer sequences (5’→3’) | |
|---|---|---|
| SPINK1 | F: AGAGGCCAAATGTTACAATG | R: ATGGGATTTCAAAACCTTGG |
| GAPDH | F: TGCACCACCAACTGCTTAGC | R: GGCATGGACTGTGGTCATGAG |

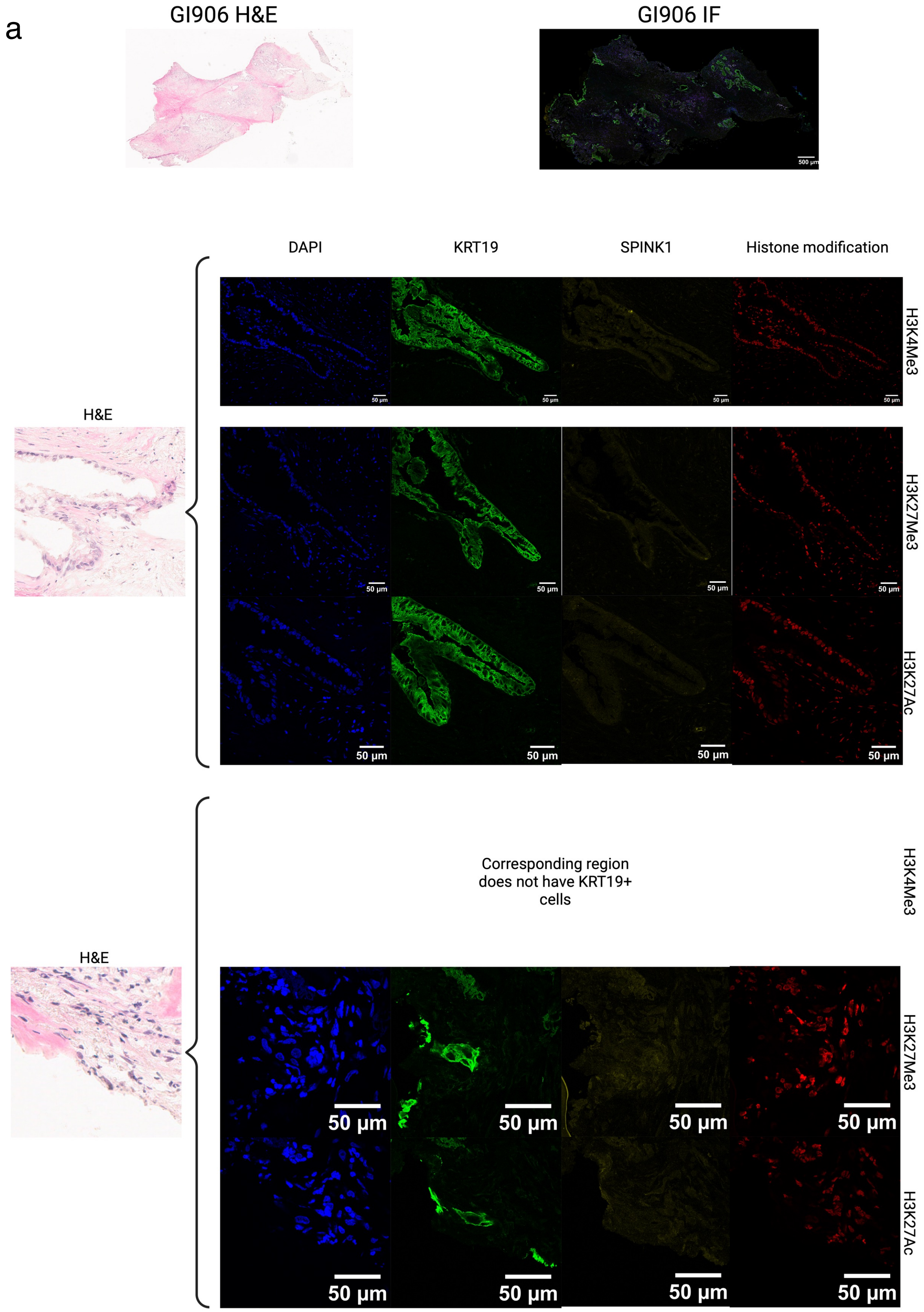
a
GI906 H&E
GI906 IF
500 µm
DAPI
KRT19
SPINK1
Histone modification
H3K4Me3
H3K27Me3
H3K27Ac
H&E
50 µm
Corresponding region
does not have KRT19+
cells
H3K4Me3
H3K27Me3
H3K27Ac
H&E
50 µm

b

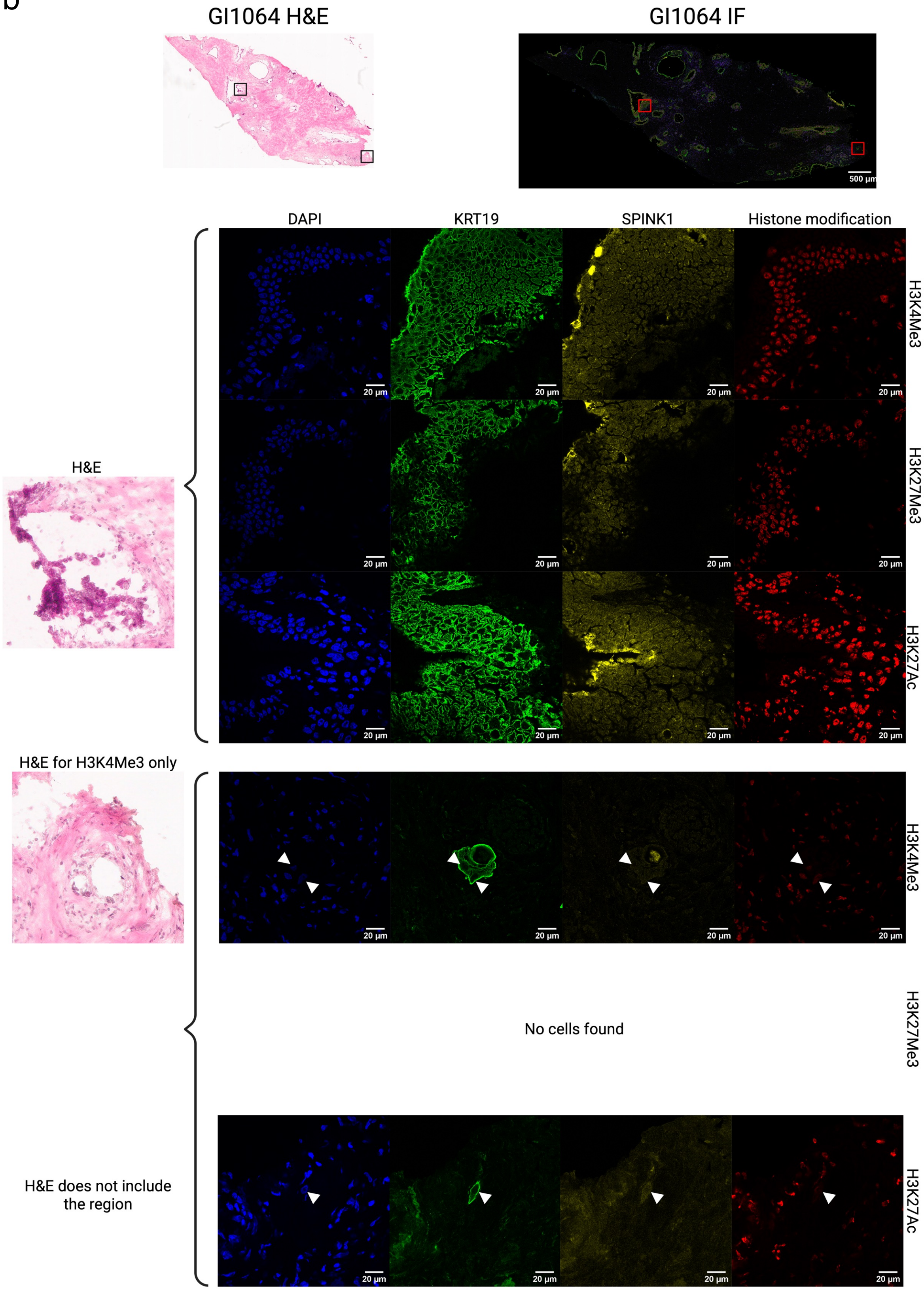
GI1064 H&E
GI1064 IF
500 µm
DAPI
KRT19
SPINK1
Histone modification
H3K4Me3
H3K27Me3
H3K27Ac
20 µm
H&E
H&E for H3K4Me3 only
No cells found
H&E does not include the region

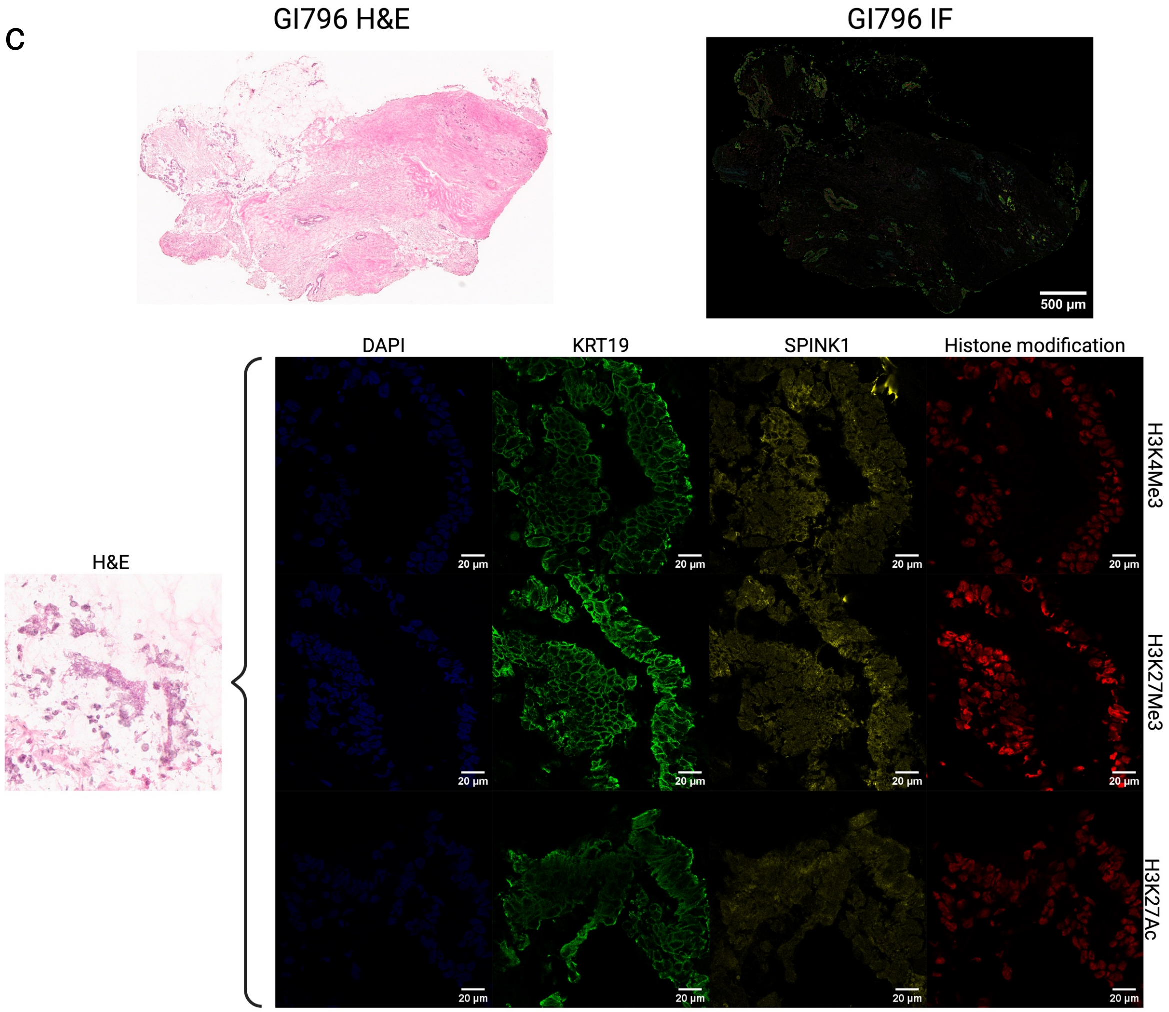


No cells found with low SPINK1 and low histone modifications

Supplementary Figure 6. H&E staining and immunostaining of DAPI (blue), KRT19 (green), SPINK1 (yellow) and histone H3 modification markers (red) for fresh frozen samples from three PDAC patients.

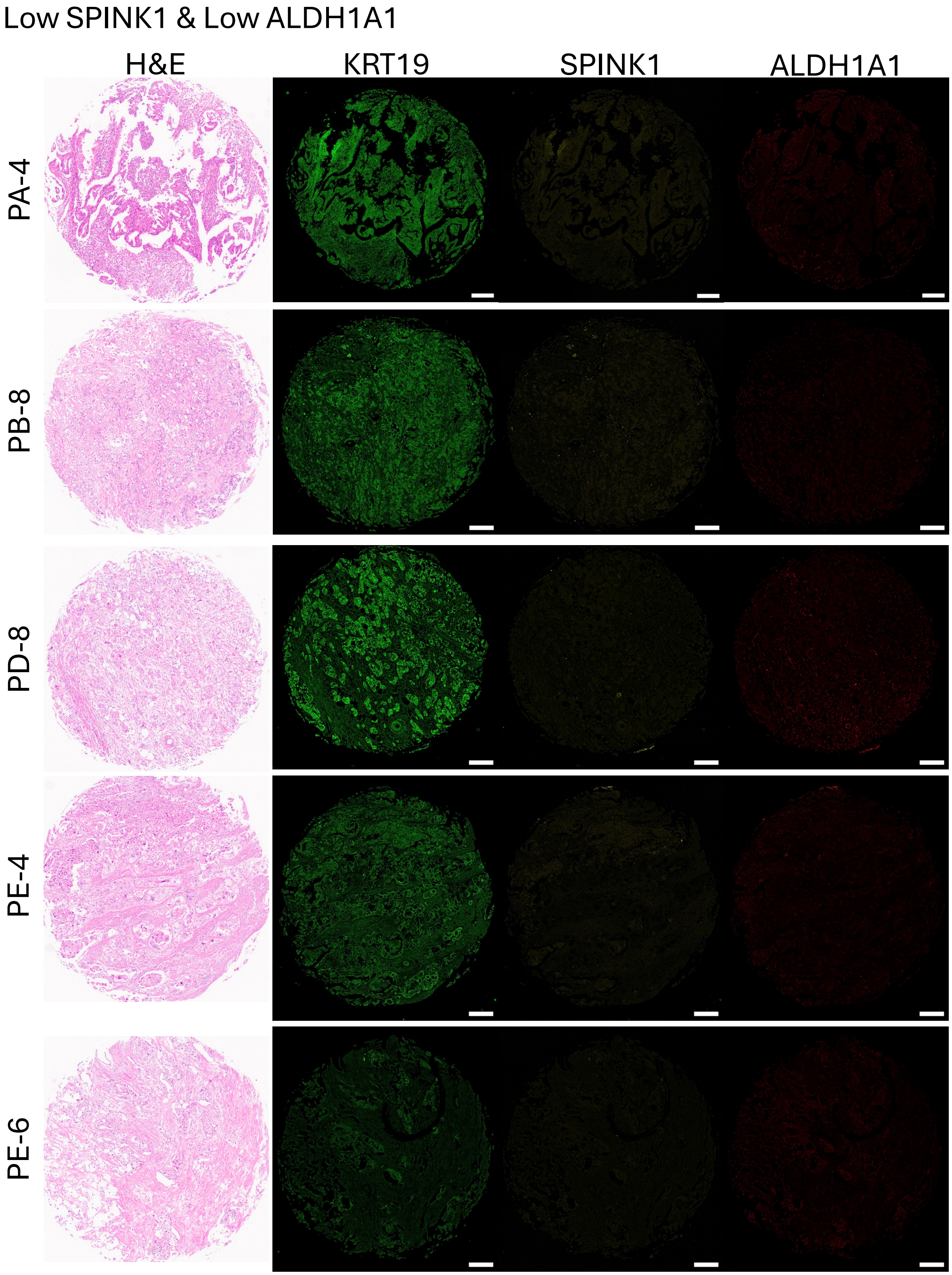
Low SPINK1 & Low ALDH1A1
H&E
KRT19
SPINK1
ALDH1A1
PA-4
PB-8
PD-8
PE-4
PE-6

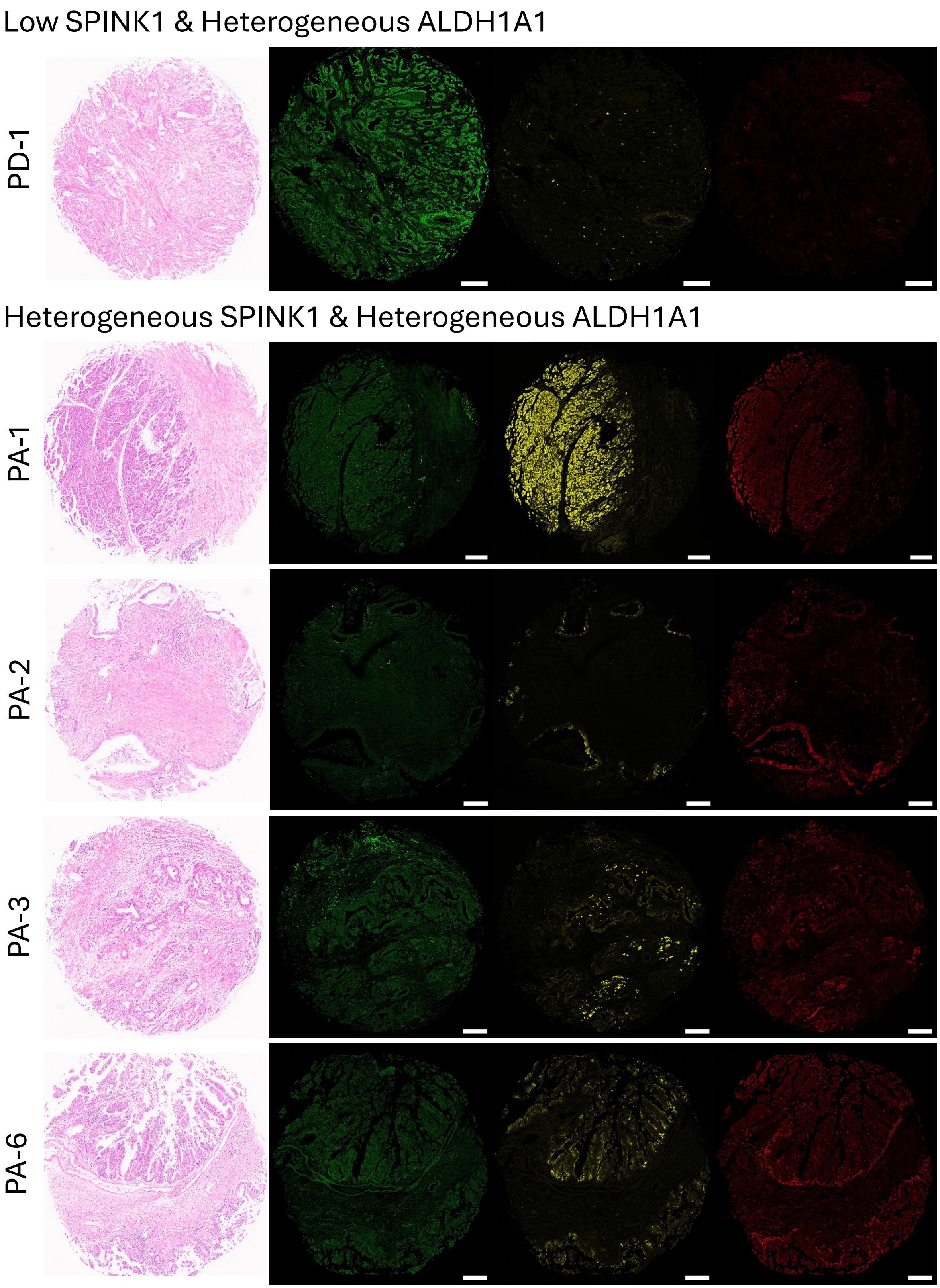
Low SPINK1 & Heterogeneous ALDH1A1
PD-1
Heterogeneous SPINK1 & Heterogeneous ALDH1A1
PA-1
PA-2
PA-3
PA-6

Heterogeneous SPINK1 & Heterogeneous ALDH1A1 (continued)

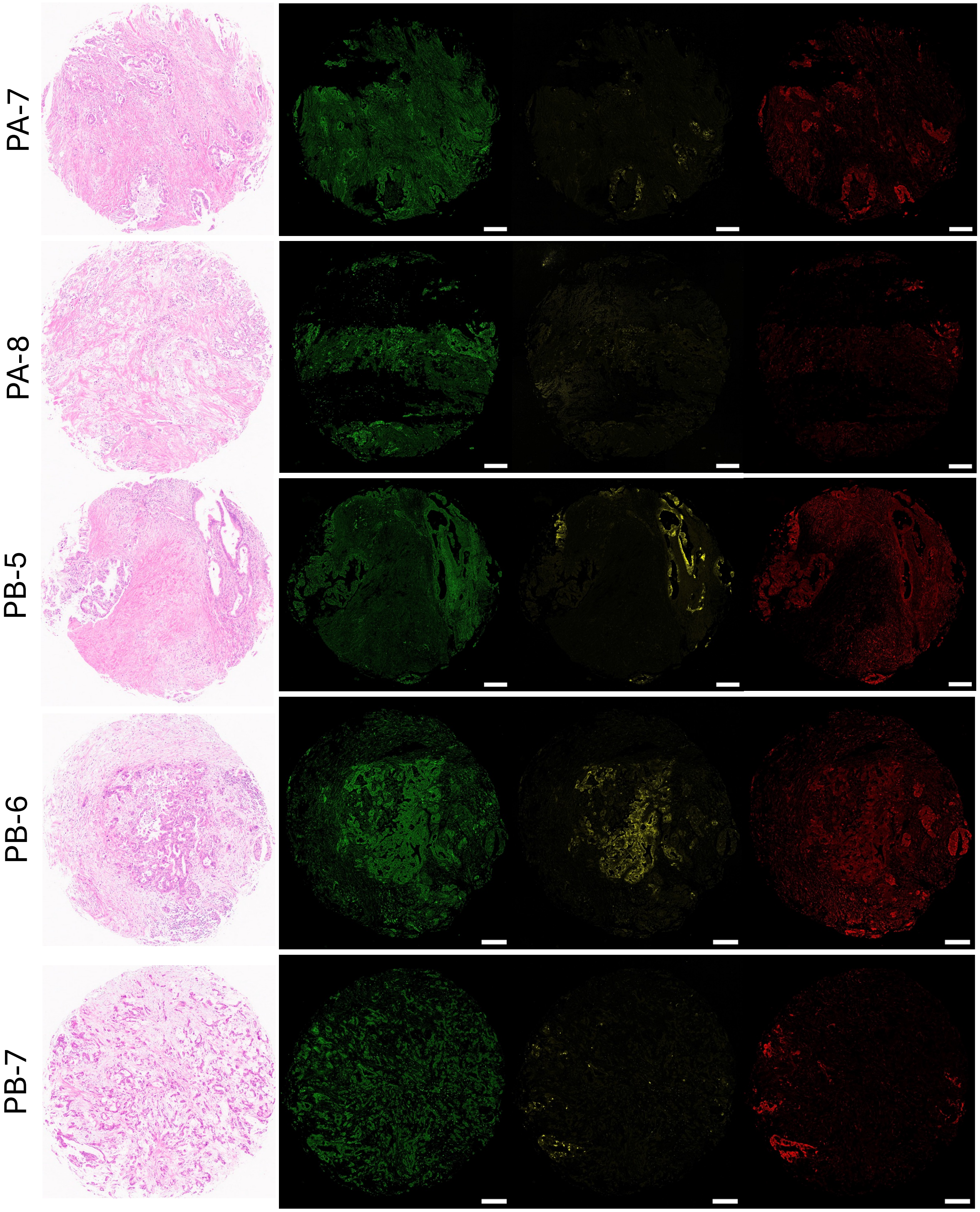

Heterogeneous SPINK1 & Heterogeneous ALDH1A1 (continued)

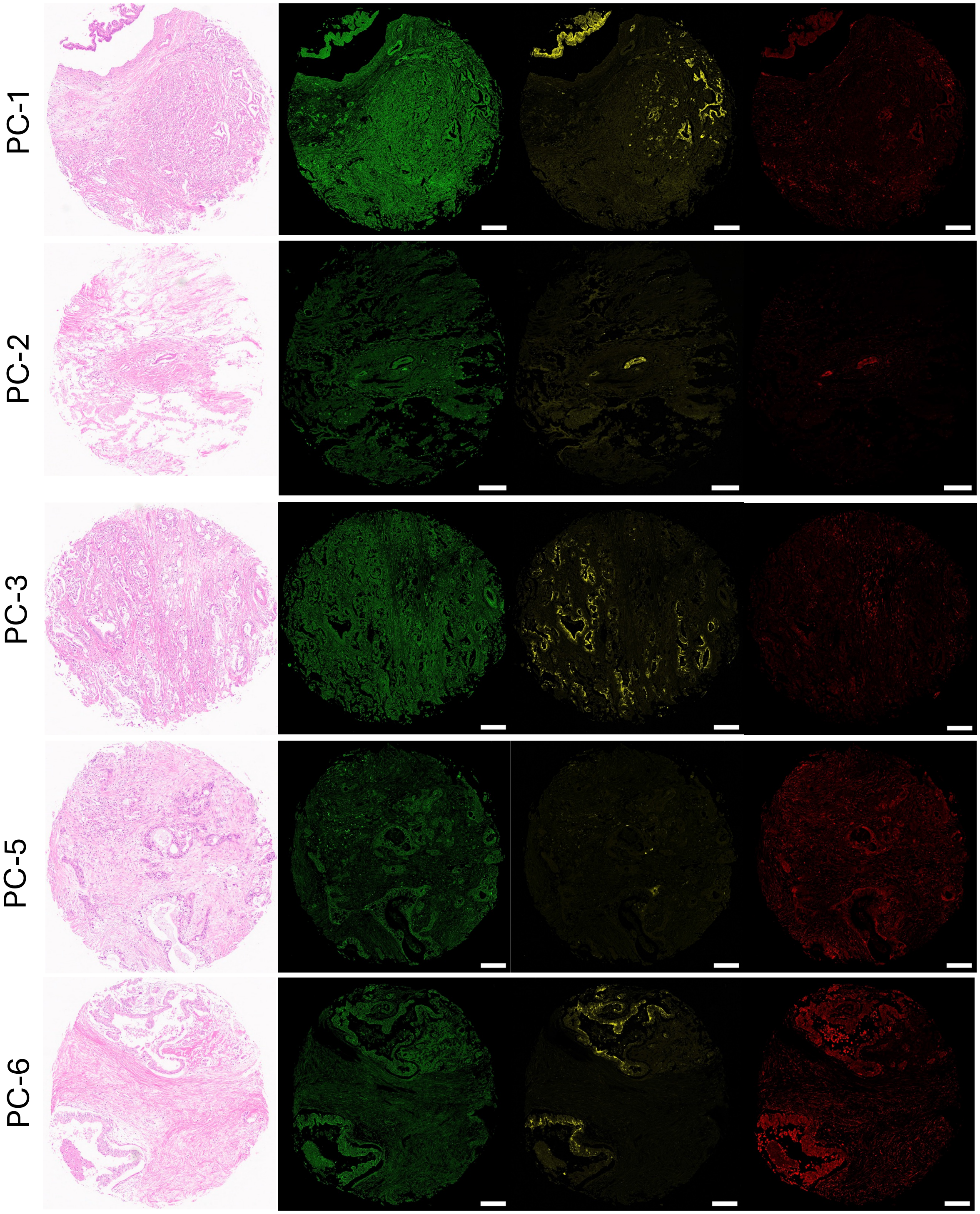

Heterogeneous SPINK1 & Heterogeneous ALDH1A1 (continued)

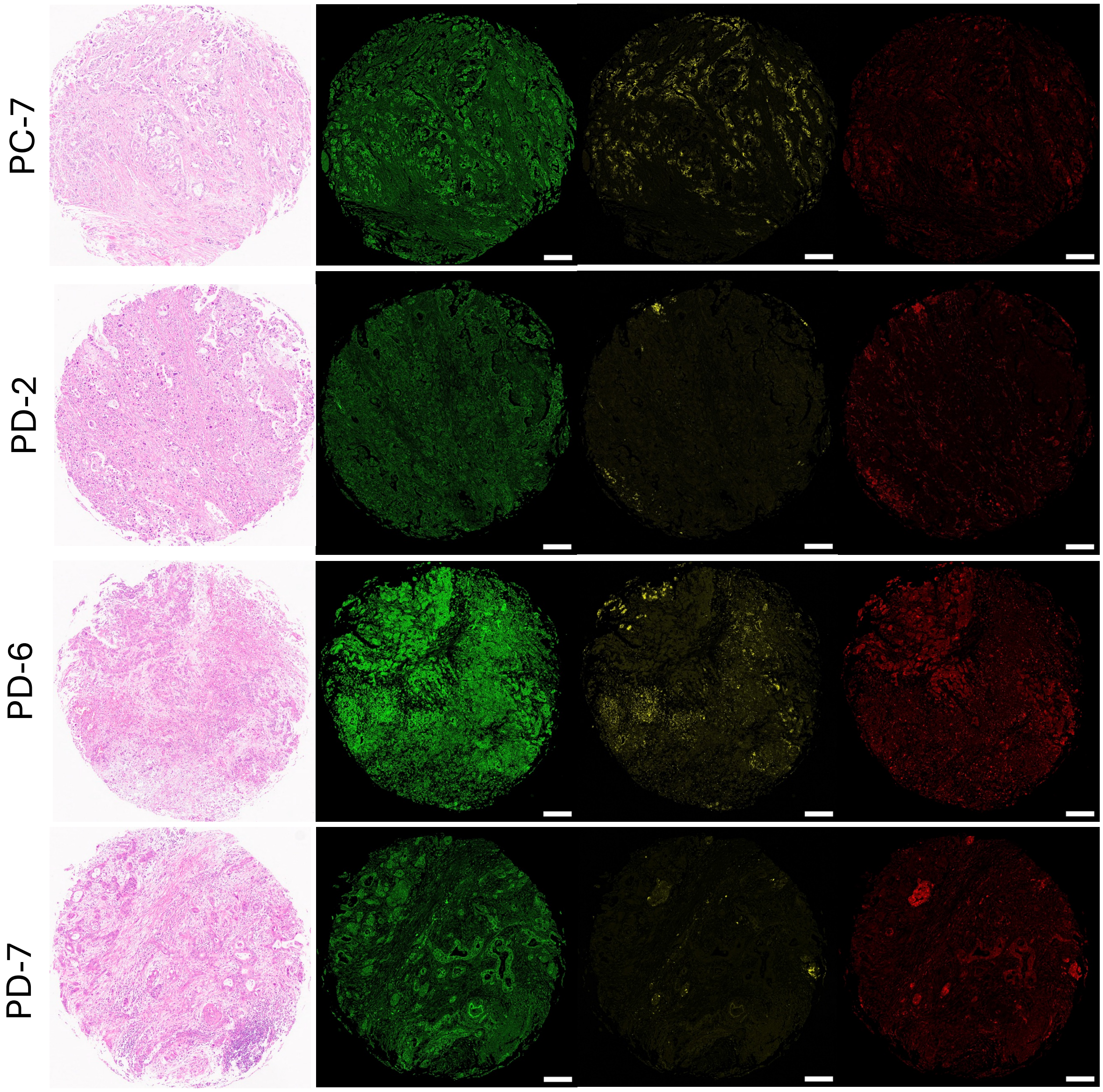

Heterogeneous SPINK1 & High ALDH1A1

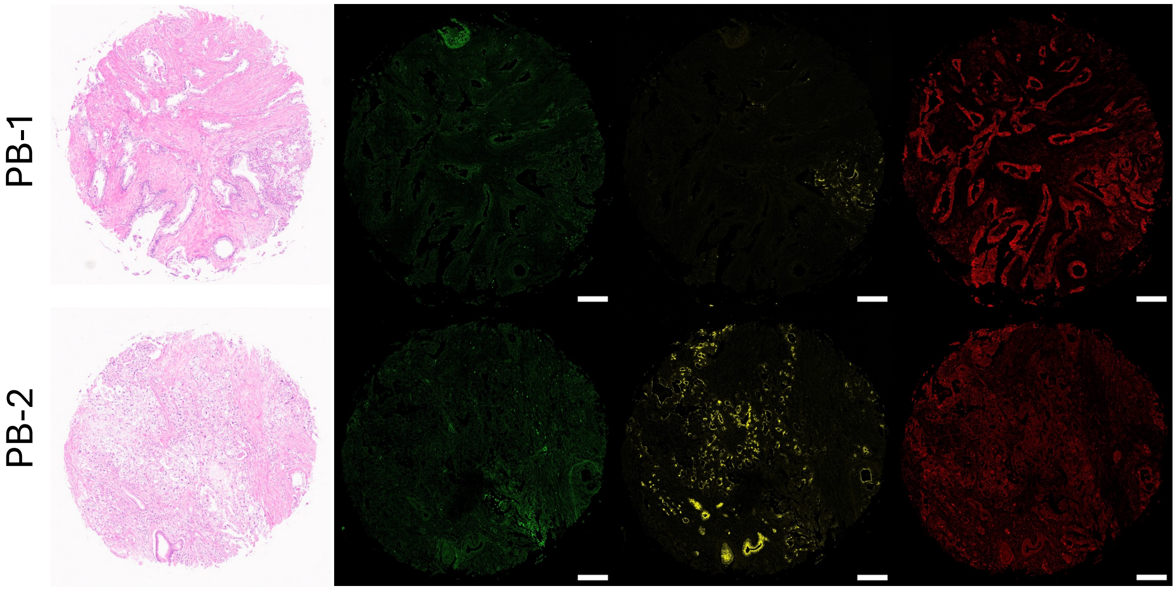


High SPINK1 & High ALDH1A1

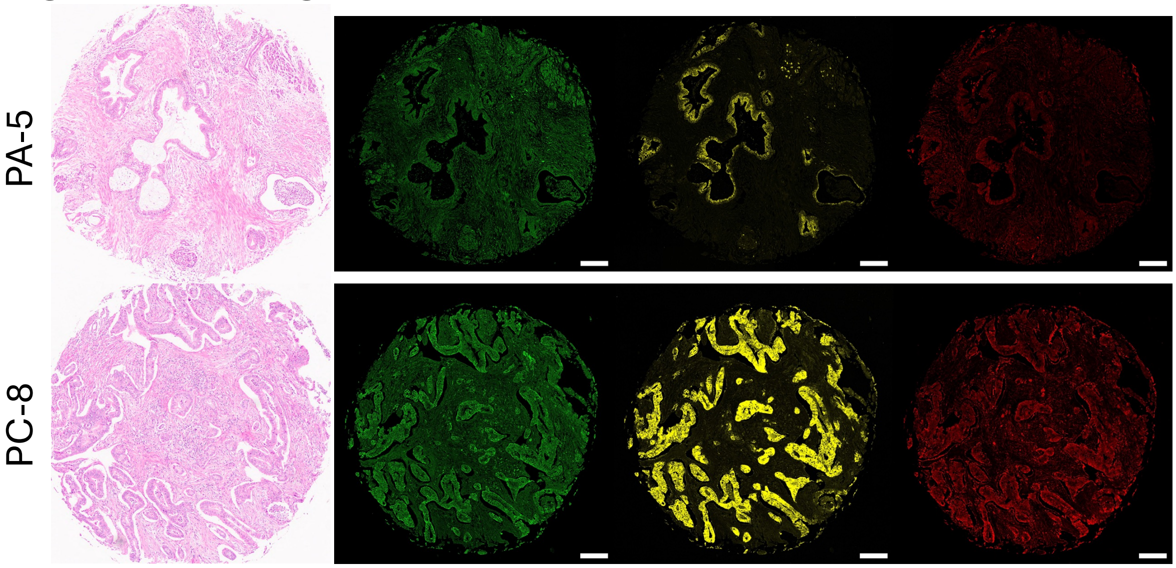


79

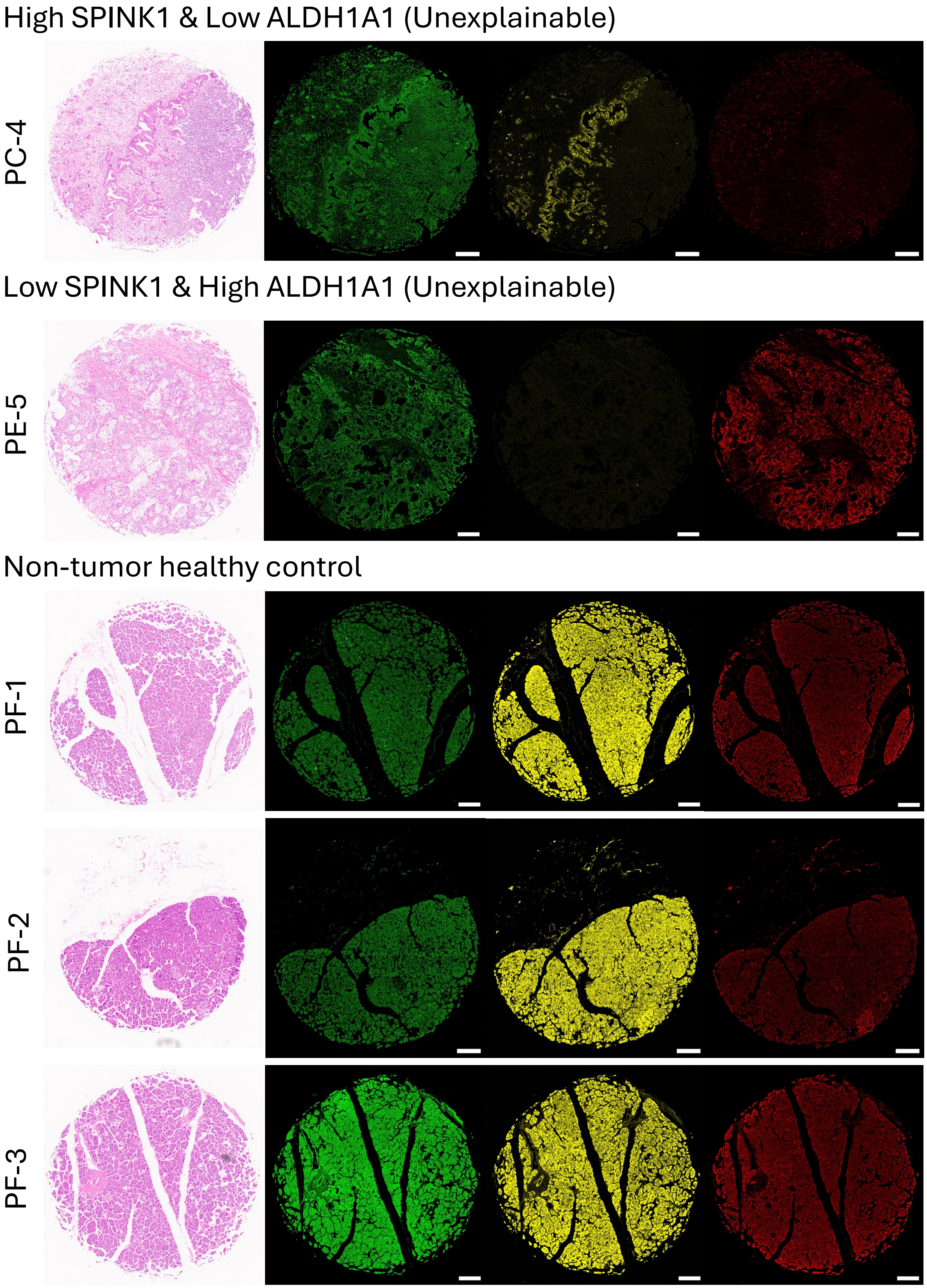
High SPINK1 & Low ALDH1A1 (Unexplainable)
PC-4
Low SPINK1 & High ALDH1A1 (Unexplainable)
PE-5
Non-tumor healthy control
PF-1
PF-2
PF-3

Non-tumor healthy control (continued)

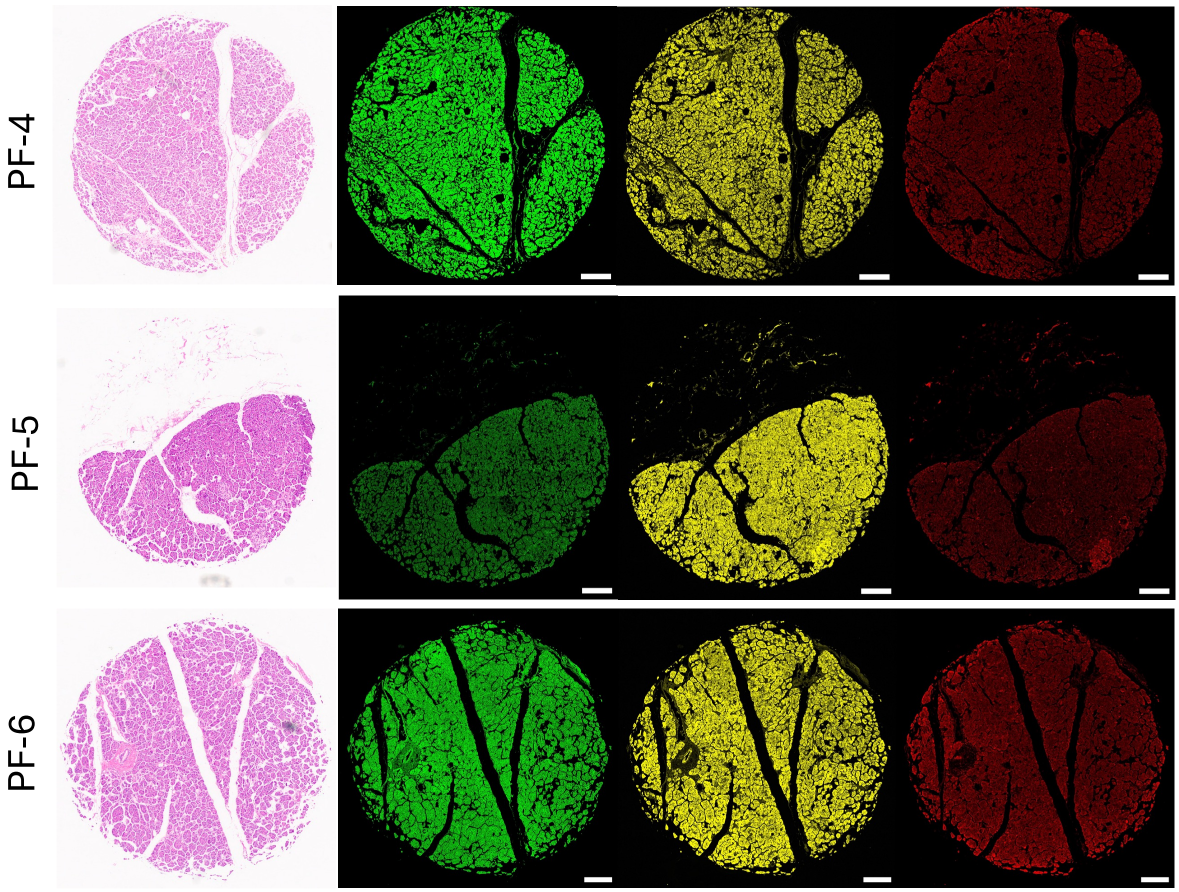


Supplementary Figure 7. H&E and immunofluorescence staining of KRT19 (green), SPINK1 (yellow), and ALDH1A1 (red) for all PDAC patient samples in TMA grouped based on the signal intensity of SPINK1 and ALDH1A1. The scale bar is 200μm.

Supplementary Table 4. The primers used for construction and sequencing of plasmids.

| Primer | Sequence (Forward) | Sequence (Reverse) |
|---|---|---|
| **SPINK1_del** | CCTAGCCGGCTCGAAGAA | CATGGTGGCGGATCCATG |
| **SPINK1_del25-34** | GAACTTAATGGATGCACC | GTCAGCTCCAGTGTTACC |
| **SPINK1_del35-44** | CCTGTCTGTGGGACTGAT | ATTGTAACATTTGGCCTCTC |
| **SPINK1_del45-54** | CCCAATGAATGCGTGTTATG | GTCATATATCTTGGTGCATC |
| **SPINK1_del55-67** | CAGACTTCTATCCTCATTC | ATAAGTATTTCCATCAGTCC |
| **SPINK1_del68-79** | CCTAGCCGGCTCGAAGAA | GCGTTTCCGATTTTCAAAACATAAC |
| **2×(GGGGS)_** UltraID | TTCAAAAATCTGGGCCTTGCGGAGGAGGTGGGAGTGGTG | TCTTCTTCGAGCCGGCTAGGGCGGTTTAAACTTAAGCTTGGTACC |

| | | |
|---|---|---|
| **SPINK1_ALFA_backbone** | CCTAGCCGGCTCGAAGAAG | GCAAGGCCCAGATTTTTGAATG |
| **SPINK1SP_UltraID** | GGAGGAGGTGGGAGTGGT | GTCAGCTCCAGTGTTACCAG |

Supplementary Table 5. The list of primary antibodies used in the Western blot studies.

| Target Name | Company | Cat # | Dilution ratio |
|---|---|---|---|
| GAPDH | santa cruz | sc-32233 | 1:1000 |
| SPINK1 | R&D systems | MAB7496 | 1:500 |
| ACTB | Cell Signaling Technology | 3700 | 1:1000 |
| CDH1 | BD biosciences | 610181 | 1:1000 |
| CDH2 | Cell Signaling | 4061S | 1:1000 |
| ZEB1 | Proteintech | 21544-1-AP | 1:1000 |
| VIM | Abcam | ab92547 | 1:1000 |
| Histone H3 | Proteintech | 17168-1-AP | 1:500 |
| H3K4me3 | Thermo Fisher | MA5-11199 | 1:500 |
| H3K27me3 | Thermo Fisher | MA5-11198 | 1:500 |
| H3K27Ac | Abcam | ab4729 | 1:500 |
| VEGFR | Cell Signaling | 2479S | 1:500 |
| p-VEGFR | Cell Signaling | 2471S | 1:500 |
| $\beta$-Catenin | Cell Signaling | 8814S | 1:1000 |
| SRC | Thermo Fisher | 701396 | 1:500 |
| p-Src Thr416 | Cell Signaling | 6943S | 1:1000 |
| NCL | Novus | NB600-241SS | 1:1000 |
| p-NCL Thr76 | Novus | NBP3-21607 | 1:1000 |
| p-NCL Thr84 | Novus | NBP3-21606 | 1:1000 |
| COL18A1 | Abcam | ab275390 | 1:1000 |
| Endostatin | Thermo Fisher | # PA1-601 | 1:1000 |
| NUP153 | Proteintech | 14189-1-AP | 1:1000 |
| CNX | Proteintech | 66903-1-Ig | 1:1000 |
| Histone H2B | Proteintech | 15857-1-AP | 1:1000 |
| Biotin | Thermo Fisher | 21130 | 1:1000 |
| ALFA-tag | nanotag biotechnologies | N1583 | 1:1000 |

| | | | |
|---|---|---|---|
| EGFR | Cell Signaling | 2232S | 1:1000 |
| p-EGFR | Cell Signaling | 4407S | 1:1000 |
